# Lithic bacterial communities: ecological aspects focusing on *Tintenstrich communities*

**DOI:** 10.1101/2024.04.30.591805

**Authors:** Francesca Pittino, Sabine Fink, Juliana Oliveira, Elisabeth M.-L. Janssen, Christoph Scheidegger

## Abstract

*Tintenstrich* communities (*TC*) are mainly composed by Cyanobacteria developing on the rock substrate and forming physical structures strictly connected to the rock itself. Endolithic and epilithic bacterial communities are important because they contribute to nutrients release within run-off waters flowing on the rock surface. Despite them being ubiquitous, little information about their ecology and main characteristics is available. In this paper, we characterized the bacterial communities of rock surfaces of *TC* in Switzerland through Illumina sequencing and investigated their bacterial community composition on two substrate types (silicious and limestone rocks) through multivariate models. Our results show that Cyanobacteria and Proteobacteria are the predominant phyla in this environment. Bacterial alpha diversity was higher on limestone than on siliceous rock, and beta diversity of siliceous rock varied with changes in rock surface structure. Here we provide novel insights into the bacterial community composition of *TC*, their differences from other lithic communities, and the effects of the rock substrate and structure.

## INTRODUCTION

We are aware that bacteria can colonize many different substrates, even in the most extreme locations (e.g., glaciers, deserts, deep oceans)^[1–3]^. Bare rocks can be considered as an extreme environment being oligotrophic, frequently desiccating and presenting high-temperature shifts. Lithic habitats also present the oldest terrestrial surface, but knowledge about life on rock surfaces is still very limited^[4]^. While bacteria are the main colonizers of this environment, archaea, algae, fungi and lichens also play an important role for the community composition^[4]^. Although these organisms are microscopically small, they form conspicuous biofilms named *Tintenstrich* (from German “ink stripes”) communities (*TC*), named after the dark color change of the rock surface. These particular and characteristic structures develop on rock surfaces with intermittent, rare to frequent, water runoff (Fig. 1a-b) ^[5]^. *TC* are semi-aquatic communities that develop on different types of rock. They are mainly composed of free-living Cyanobacteria, and cyano-lichens can also occur^[4]^. They can be recognized thanks to the change in characteristic rock color, while the surface 3D morphology in the presence and absence of *TC* remains largely unchanged. Typically, *TC* form extended, contiguous communities that are smooth as the bare rock and often become slippery when wet because of characteristic gelatinous sheaths of cyanobacterial cells. In other cases *TC* can also be fragmented and scattered among other structures such as bryophytes and epilithic lichens that are associated with green algae or Cyanobacteria (Figure 1).

**Figure 1.**
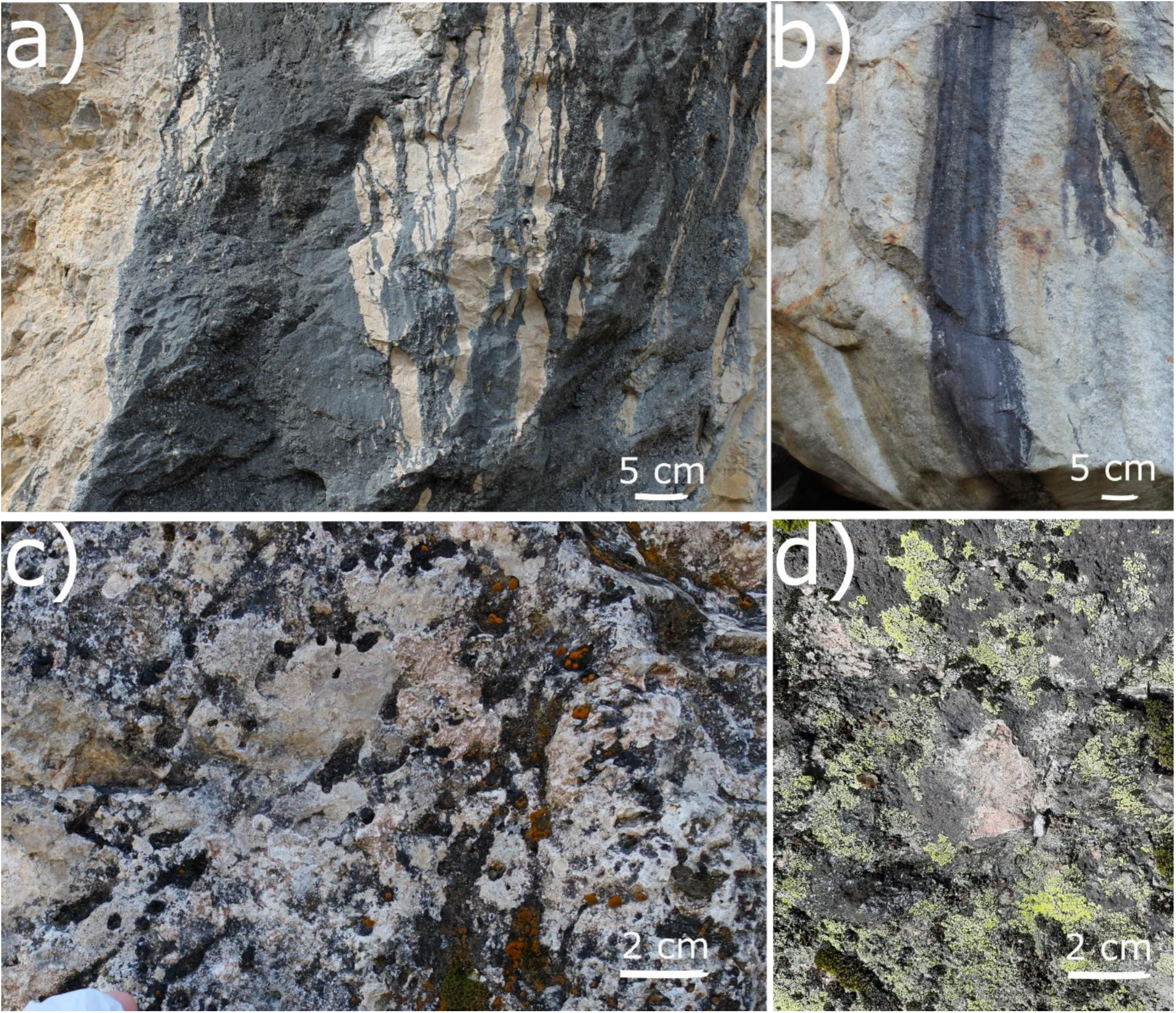
Examples of contiguous TC on limestone (a) and siliceous rock (b). Examples of fragmented TC scattered among green-algal lichen communities on limestone (c) and siliceous rock (d).

Many environmental variables can affect *TC*, but so far the main known properties recognized promoting their development are the water retention capability of the rock^[4]^, sun exposure^[6]^ and the type of rock^[7]^. *TC* are widely occurring in both anthropic and natural environments since they can form on sculptures and architectural works^[8]^ and on newly exposed rocks, for example, after glaciers retreat, which is now occurring at unprecedented speed^[9]^. *TC* also play an important role in rock weathering and biogeochemical cycles both directly on the rock and “down-valley” where inorganic nutrients are released as a consequence of bio-weathering through water flowing across *TC* surfaces^[4,10]^.

The relationship established between the bacterial community colonizing the rock and the substrate itself is particularly strong. Three categories of bacterial communities can be recognized, according to which part of the substrate is colonized: (i) epilithic bacteria, which stay on the external surface, (ii) hypolithic bacteria, which stay on the ventral surface of rocks (i.e., beneath pebbles), and (iii) endolithic bacteria, which develop inside the rock, typically few millimeters deep, so that light can still penetrate to allow for photosynthesis. The endolithic communities can then occupy different microhabitat inside the rock itself and, according to which one they inhabit, they can be further categorized as: chasmoendoliths (inhabiting fissures and cracks), cryptoendoliths (occurring in preexisting cavities and pores) and euendoliths (which actively penetrate and create their habitat into the rock substrate)^[4]^. These microenvironments allow the instauration of a certain variability on an apparently homogeneous rock substrate. Therefore, the micro-architecture of the rock drives the different taxa to colonize different parts according to their metabolisms and characteristics^[4,11]^. Versatile metabolisms and resistance to high stresses are good characteristics that select for certain strains in this kind of habitat. The most abundant bacterial phyla described in endolithic communities are: Cyanobacteria, Actinobacteria and Proteobacteria^[12]^. They are all good colonizers of different types of rocks and they can survive exposed to different environmental conditions in both aquatic and terrestrial environments^[12]^. These communities fulfill important ecological roles due to their different and versatile metabolisms as they can fix nitrogen, complete the sulfur cycle and release acids that decrease pH and provoke minerals dissolution^[12]^, and they are also likely responsible for the soil formation^[13]^.

Cyanobacteria are good colonizers of bare rock surface as they are phototrophic, they can fix carbon and some genera are also capable of nitrogen fixation, they have pigments protecting them from high UV radiation and, in the most extreme conditions, they can enter a dormant state^[14]^. Indeed, they are a photoautotrophic bacterial phylum able to colonize different extreme environments and, because of their metabolism, they can also provide organic carbon and nitrogen for heterotrophic taxa^[15]^. Another important characteristic, typical of both bacteria and fungi to survive on rock surfaces, is the ability to produce extracellular polymeric substances (EPS). These compounds provide protection against desiccation and, thanks to their adhesive properties, can retain nutrients but they also promote biodeterioration^[4]^. Most of the studies on Cyanobacteria conducted so far focused on toxin producing blooms in aquatic environments ^[16]^, on soil crusts because of their contribution to biogeochemical cycles through nitrogen and carbon fixation^[17]^, and on glaciers since Cyanobacteria are good pioneer colonizers of these environments^[18]^. The importance of Cyanobacteria on rock surfaces, such as *TC,* has still been widely overlooked. In addition, other organisms (e.g. gasteropoda) can spend part of their life on rock substrate, actively feeding on cyanobacteria and cyano-lichens of *TC* and hence directly connecting them to the whole trophic network^[19]^.

Previous studies investigating bacterial communities on rocks surface used both molecular and microscopy approaches, and examined both small areas or more extended ones within the same geographic regions^[6,12,20–22]^. Therefore, some information is now available about bacterial communities of rock with a special focus on Cyanobacteria, while knowledge on *TC* is missing. The ecology of *TC* was first described in 1945 by the Swiss biologist Otto Jaag^[23]^ and Lüttge (1997)^[5]^, but no further information exists about these important structures.

Therefore, in this article, we aim to answer to two three main questions:

- Which are the main factors driving lithic bacterial community composition and alpha diversity?
- Are differences related to the rock substrate composition, light exposition or elevation?
- Do *TC* composition differ from other rock communities?

## MATERIALS AND METHODS

### Sampling design and samples collection

A total of 209 rock samples were collected in Switzerland in 19 sampling areas (Fig. 2, Tab. S1) with a sterilized hammer and chisel. At least three replicates were collected per study sites. Sampling areas were selected to obtain a similar sample number of both limestone and siliceous rock. Some of the sampling areas were previously reported in literature on endolithic bacteria^[22,23]^.

**Figure 2.**
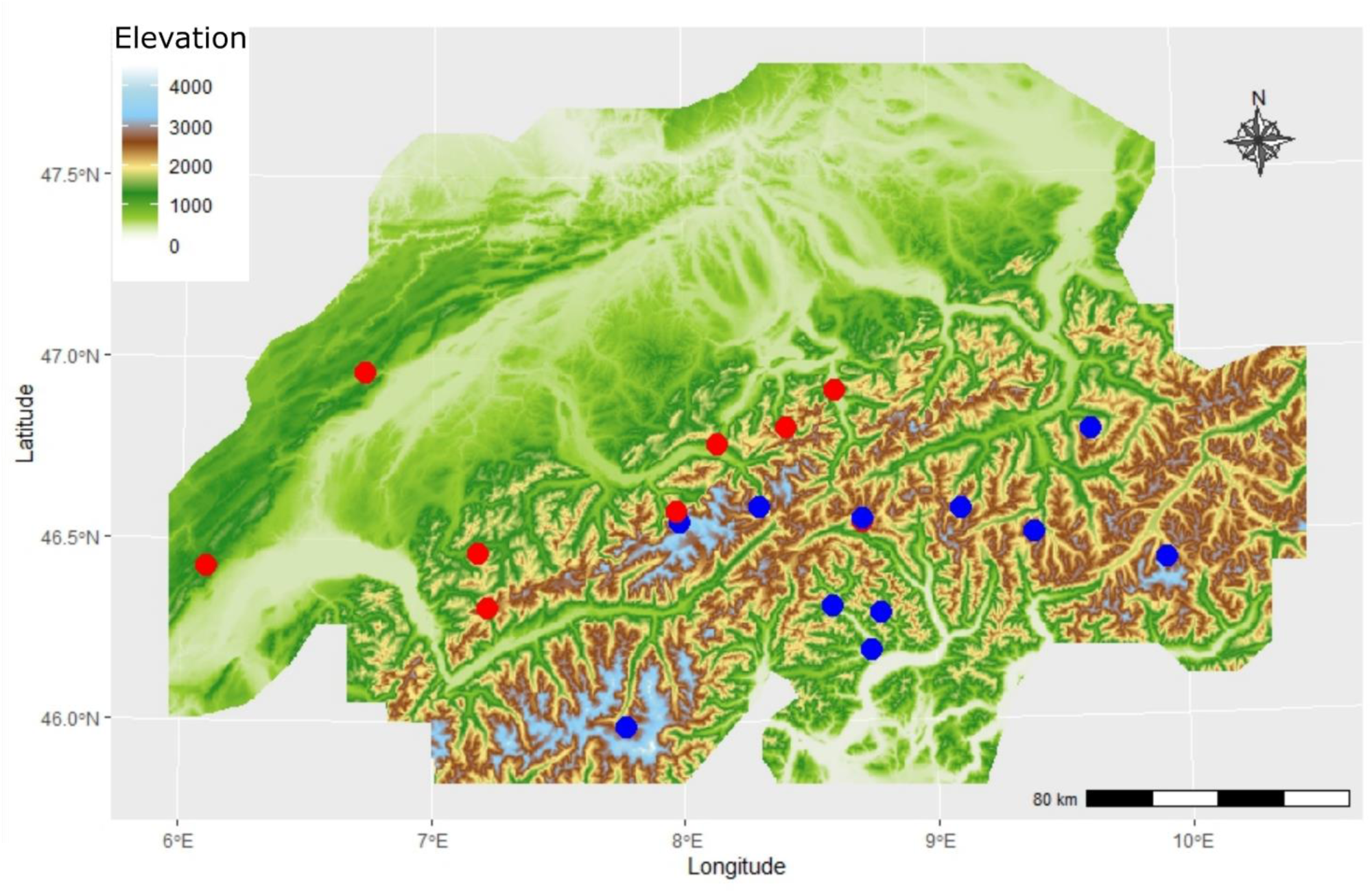
Map of 19 sampling areas in the Swiss Alpine region at different elevations (color bar), different substrate of either limestone (blue circles, n=12) or siliceous rock (red circles, n=9). Replicate were taken with different light exposition (north and south facing, not shown) adding up to a total of 92 limestone samples and 117 siliceous rock samples.

Samples were collected removing the rock, preferably with contiguous smooth *TC* when present (Fig. 1a-b, referred to as contiguous *TC*). If contiguous *TC* were not available, rock samples with fragmented *TC* scattered among bryophytes and epilithic lichens were collected (Fig. 1c-d, referred to as fragmented *TC*). Exposition, elevation and fragmentation were registered. Fragmentation is used as a factor variable, which indicates how and if the rock presents a classic contiguous *TC.* In details, we categorized fragmentation as a three levels variable: Level 1 being contiguous *TC* with the same 3D morphology of the rock itself and no evident growth of green-algal lichens and bryophytes (Fig. 1a-b), level 2 where *TC* were highly fragmented and scattered among the dominating green-algal lichens and bryophytes (Fig. 1c-d), and Level 3 as a condition in-between the previous two where *TC* formed a macroscopically visible mosaic with bryophytes and epilithic lichens. Samples were stored at −20 °C until processing.

### DNA extraction and sequencing

DNA was extracted from rock pieces scratched from the surface (approx. from the first 3-4 mm) with the help of a sterile chisel and of a handheld rotary power tool (Dremel 8220-1, Bosch NL). DNA extraction was performed from 0.25 g of lyophilized scratched rock with the DNeasy PowerSoil Pro Kit (QIAGEN), using two metal beads in a sterile 2 mL Eppendorf instead of the PowerBead Pro Tube, and adding a 2 minutes vortexing in dry conditions as first step. The other steps were performed according to the manufacturer’s instruction (protocol dated May 2019). A first PCR was performed on the V4-V5 hypervariable region of the 16S rRNA to check for DNA quality and inhibition using the original DNA and both 1:10 and 1:100 dilutions. PCR mixture was composed of 7.5 µL of JumpStart™ REDTaq® ReadyMix™ (Sigma-Aldrich), 0.75 µL of each primer 10 µM (515F and R926^[24]^), 4.5 µL of Milliq water and 1.5 µL of DNA. The PCR program was the following: 3 minutes of initial denaturation at 95°C, 28 cycles of 45 seconds at 95°C, 45 seconds at 50°C, and 90 seconds at 72°C, and a final extension of 5 minutes at 68°C. A second PCR was performed with the KAPA HiFi Hotstart ReadyMix (Roche) and 10 µM of the two primers 515F and 806R to amplify the V4 hypervariable region of the 16S rRNA, for a final volume of 2 x 20 µL per sample. Primers were modified with Illumina adapters, a shift and a linker according to Kozich et al. ^[25]^ for the 515F and according to Caporaso et al. ^[26]^ for the 806R. The PCR program was the following: 3 minutes of initial denaturation at 95°C, 28 cycles of 45 seconds at 95°C, 45 seconds at 58°C, and 45 seconds at 72°C, and a final extension of 5 minutes at 72°C. PCR products were sent to the NGS Platform of the Institute of Genetics at Bern University (Bern, Switzerland), for sequencing with MiSeq Illumina platform (Illumina, Inc., San Diego, CA) using a 2 × 250 bp paired-end protocol.

### Bioinformatic and statistical analyses

Demultiplexed reads were clustered in Amplicon Sequences Variants (ASVs) with DADA2^[27]^. ASVs were taxonomically classified using the SILVA database^[28]^. Singletons (ASVs present in one sample only) were removed from the database to avoid inflation of the variance explained by multivariate tests^[29]^.

Alpha-diversity was investigated trough the number of ASVs, the Shannon diversity index^[30]^ and the Gini inequality index^[31]^, which were calculated on a dataset rarefied to 4155 sequences per sample, this number is slightly less than the lowest number of sequences in a sample.

Beta-diversity was investigated on a non-rarefacted dataset where ASVs numbers were transformed with the Hellinger distance^[32]^. Beta-diversity was investigated through the Canonical Correspondence Analysis (CCA) of the ASVs abundances transformed with the chi-square distance using as predictors northness (cos(aspect)), eastness (sin(aspect)), elevation and the interaction between rock type (limestone or siliceous rock) and fragmentation.

Post-hoc tests (Tukey method) were used to assess pairwise differences in the structure of bacterial communities between rock and levels of fragmentation, correcting P-values with the false discovery rate (FDR) method using the Benjamini–Yekutieli procedure ^[33]^. Variation partitioning was used to quantify the variation of community structures according to the same variables used in the CCA. The most abundant phyla and orders were also investigated through generalized linear models (GLMs) with a Poisson distribution corrected for overdispersion and correcting P-values using the same FDR procedure as above. The most abundant taxa were considered those that, when summed up, represent together more than 80% of the whole community ASVs. The multipatt procedure was used on the most abundant genera to investigate indicator species in *TC* correcting P-values with the false discovery rate (FDR) method using the Benjamini–Yekutieli procedure. Indicator genera were investigated taking into account all the ecologically significant groups of the three levels of the variable level of fragmentation. We considered as ecologically significant all the levels of fragmentation in combinations with the rock type. These levels can fit along a fragmentation gradient.

Shared ASVs were investigated to better describe the bacterial communities of the samples collected from contiguous *TC* of limestone and siliceous rock respectively merging samples coming from the same rock wall. Analyses were performed with R 4.2.1 ^[34]^ with the VEGAN, BIODIVERSITYR, MULTTEST, INDICSPECIES and MULTCOMP packages. Finally, shared ASVs among contiguous limestone *TC* samples and among contiguous siliceous rock samples were investigated. For this aspect, contiguous *TC* samples were considered only following the conservative assumption that they represent *TC* in a strict sense and should therefore more likely provide data about the core community of these structures without too much noise. Some ASVs that resulted interesting indicator genera, were subsequently blasted on *ncbi* database (https://www.ncbi.nlm.nih.gov/) to have a better classification at genus level.

## RESULTS

We collected overall 92 limestone samples and 117 siliceous rock samples for a total of 19 sampling areas. Limestone samples had an elevation range of 500-2930 m a.s.l. with a median of 2000 m a.s.l., of which 35 samples were from contiguous *TC*, 30 samples were mainly fragmented rocks and 27 samples were both contiguous *TC* and fragmented rock surface. Siliceous rock samples had an elevation range of 370-3500 m a.s.l. with a median of 2105 m a.s.l., of which 48 samples were contiguous *TC*, 40 samples were mainly fragmented rocks and 29 samples were both contiguous *TC* and fragmented rock surface.

The number of obtained sequences per sample ranged between 4156 and 181806. The most abundant phyla in ascending order resulted to be Cyanobacteria, Proteobacteria, Actinobacteriota, Chloroflexi and Planctomycetota (Fig. 3).

**Figure 3.**
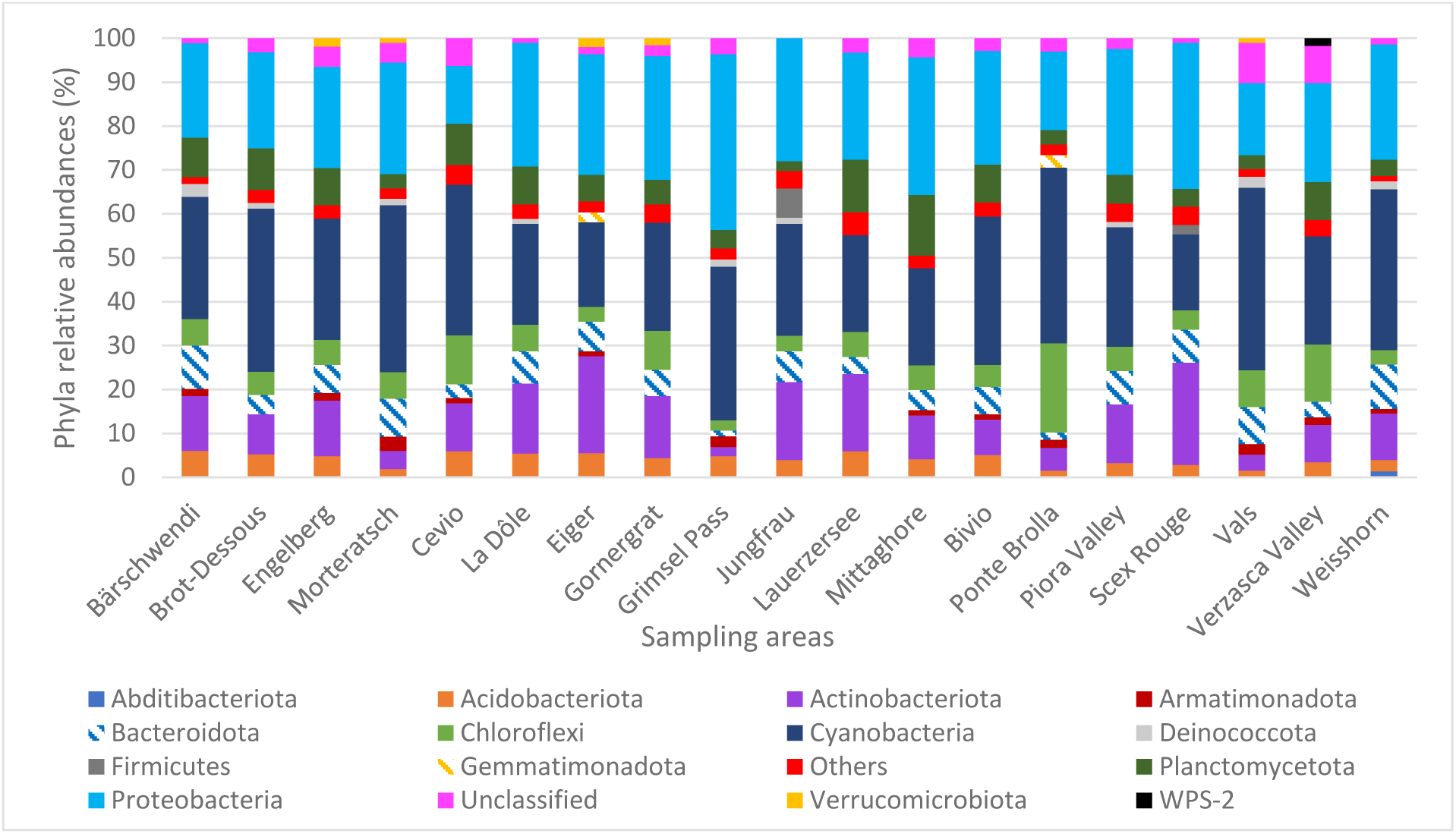
Relative abundance of bacterial phyla expressed as the percentage of sequences for the 19 sampling sites. Phyla whose abundance was <1% were grouped in ‘Others’.

At order level, the most abundant orders in ascending relative abundance resulted to be: Cyanobacteriales, Acetobacterales, Sphingomonadales, Solirubrobacterales, Rhizobiales, Isosphaerales, Cytophagales, Thermomicrobiales, Gemmatales, Blastocatellales, Ktedonobacterales, Leptolyngbyales, Burkholderiales, Caulobacterales, Rhodobacterales, Rubrobacterales, Frankiales, Pseudomonadales, Propionibacteriales, Pseudonocardiales, Chitinophagales, Armatimonadales, Deinococcales, Pyrinomonadales, Tistrellales, Tepidisphaerales, Gemmatimonadales, Chthoniobacterales, Gaiellales, Chloroflexales, Kallotenuales, Abditibacteriales, Gloeobacterales, Microtrichales, Bacillales, Thermosynechococcales, Vicinamibacterales, Euzebyales, Micrococcales and Kineosporiales (Fig. S1).

The CCA showed that all the variables included in the model were significant in explaining the variance present in the bacterial community (Tab. 1, Fig. 4, Fig. S2 shows the same plot as Fig. 4 with a focus on selected sampling sites (Piora Valley and Weisshorn) where both limestone and siliceous rock were found).

**Figure 4.**
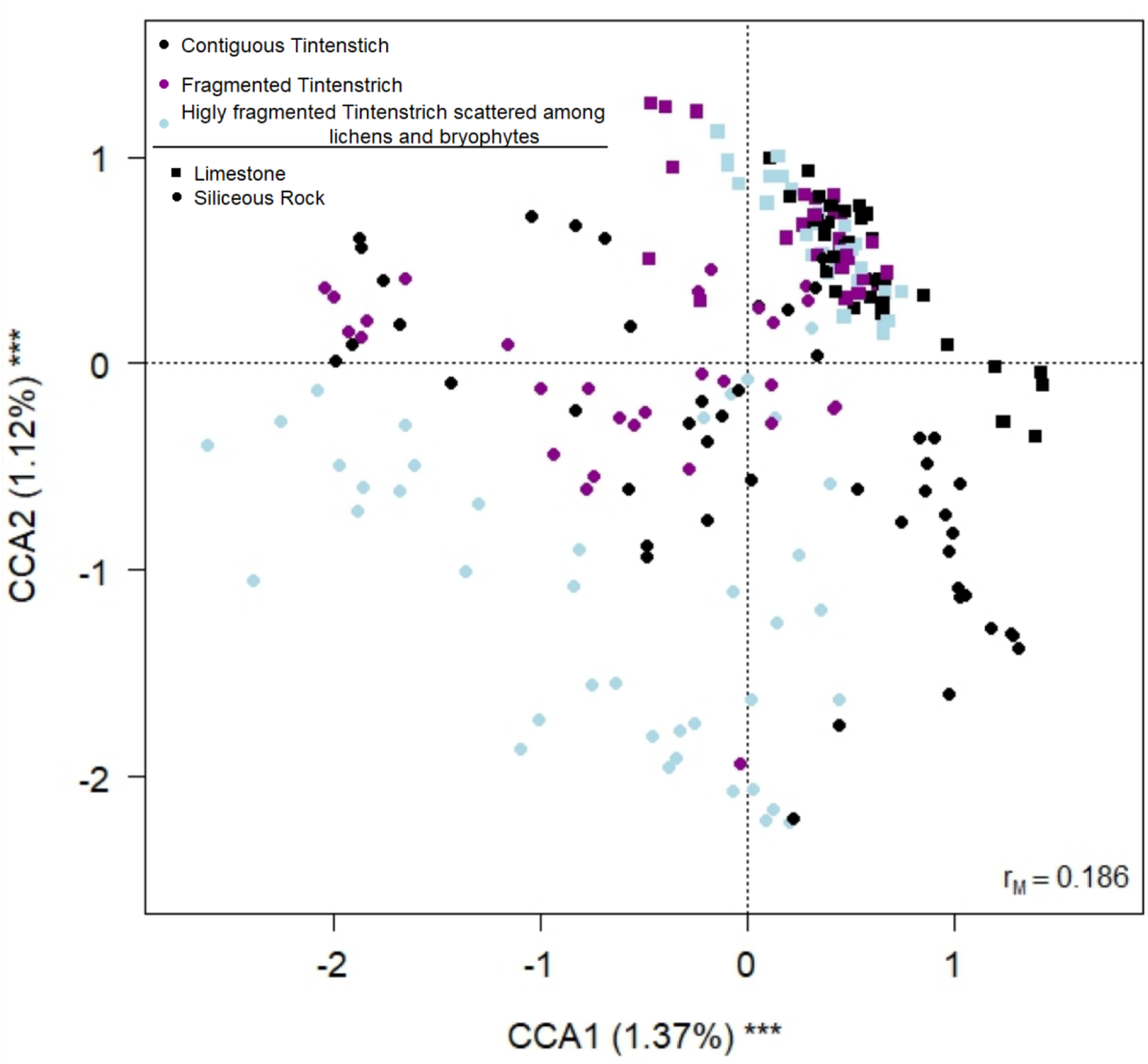
Biplot from the Canonical Correspondence Analysis on bacterial Amplicon Sequences Variants abundance on fragmentation, rock type, elevation, northness and eastness. Each data point represents one sample. The fragmentation is indicated by different colours (black = contiguous TC, purple = both contiguous TC and fragmented surface, light blue = fragmented rock surface). The arrow indicates the increase in elevation. Squares indicate limestone samples and circles siliceous rock samples. The percentage of variance explained by each axis and its significance (***: P< 0.001) is reported. r_M_ is the Mantel correlation coefficient between the chi-square distance between samples and the Euclidean distance between the corresponding symbols in the graph. Values close to one indicate that the graph correctly represents the distance between samples.

**Table 1.**
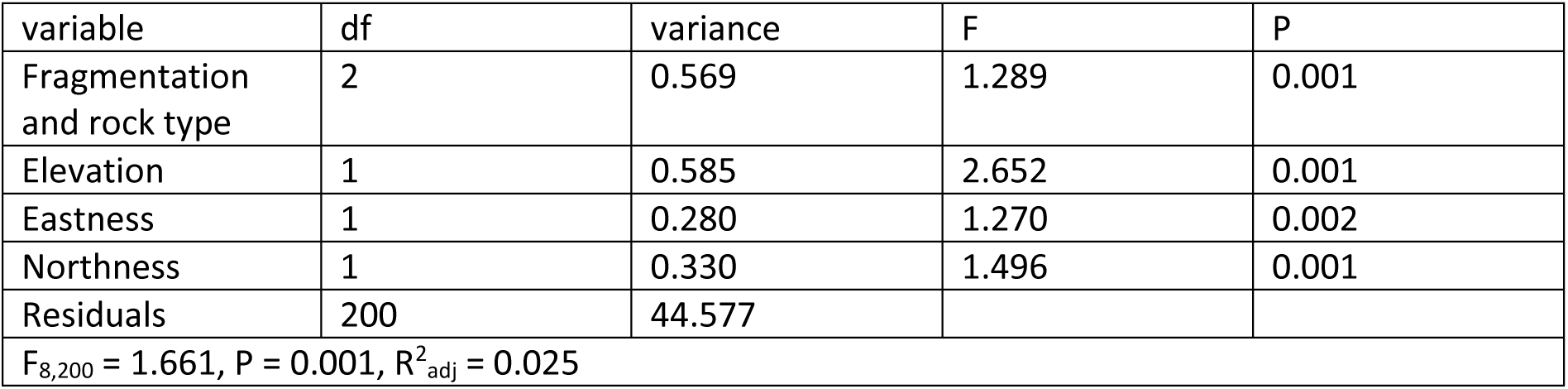
Canonical Correspondence Analysis of variance of bacterial Amplicon Sequences Variant abundance according to the interaction of fragmentation and rock type (limestone or silicious), elevation, northness, and eastness.

| variable | df | variance | F | P |
| --- | --- | --- | --- | --- |
| Fragmentation and rock type | 2 | 0.569 | 1.289 | 0.001 |
| Elevation | 1 | 0.585 | 2.652 | 0.001 |
| Eastness | 1 | 0.280 | 1.270 | 0.002 |
| Northness | 1 | 0.330 | 1.496 | 0.001 |
| Residuals | 200 | 44.577 |  |  |
| F<sub>8,200</sub> = 1.661, P = 0.001, R<sup>2</sup><sub>adj</sub> = 0.025 |  |  |  |  |

The variation partitioning revealed that, among the variables we included in the CCA model, the interaction between the type of rock and the fragmentation explained most of the variance (0.2%), also the elevation was explaining part of the variance (0.1%), while eastness and northness did not explain a part of the variance significantly different from zero. Post-hoc tests on the interaction between rock and fragmentation revealed that the bacterial community changed significantly among all the comparisons (|t_200_| ≥ 1.208, P_FDR_ ≤ 0.046) except for contiguous limestone and both contiguous limestone and fragmented rock surface (|t_200_| = 0.949, P_FDR_ = 1). In Fig. S2 we further show the same CCA of Fig. 4, evidencing four sampling areas (Weisshorn, Piora Valley, Jungfrau and Eiger). Weisshorn and Piora Valley had two different types of rocks, as well as Jungfrau and Eiger, which are distinct sampling sites yet geographically in close proximity (within 3.5 km distance).

The analyses on the alpha-diversity showed that the number of ASVs, Gini index and Shannon index did not change with the elevation (F_8,200_ ≥ 3.402, P_FDR_ = 0.0636), eastness (F_8,200_ ≥ 0.0272, P_FDR_ = 1), northness (F_8,200_ ≥ 2.920, P_FDR_ = 0.185), fragmentation level (F_8,200_ ≥ 3.038, P_FDR_ = 0.0841) and the interaction between rock and fragmentation (F_8,200_ ≥ 0.176, P_FDR_ ≥ 0.411). Significant differences were detected only among rock types (F_8,200_ ≥ 25.931, P_FDR_ < 0.001). In particular, ASV number and Shannon index were higher in limestone samples than in siliceous rock, while Gini index was lower in limestone (Fig. 5).

**Figure 5.**
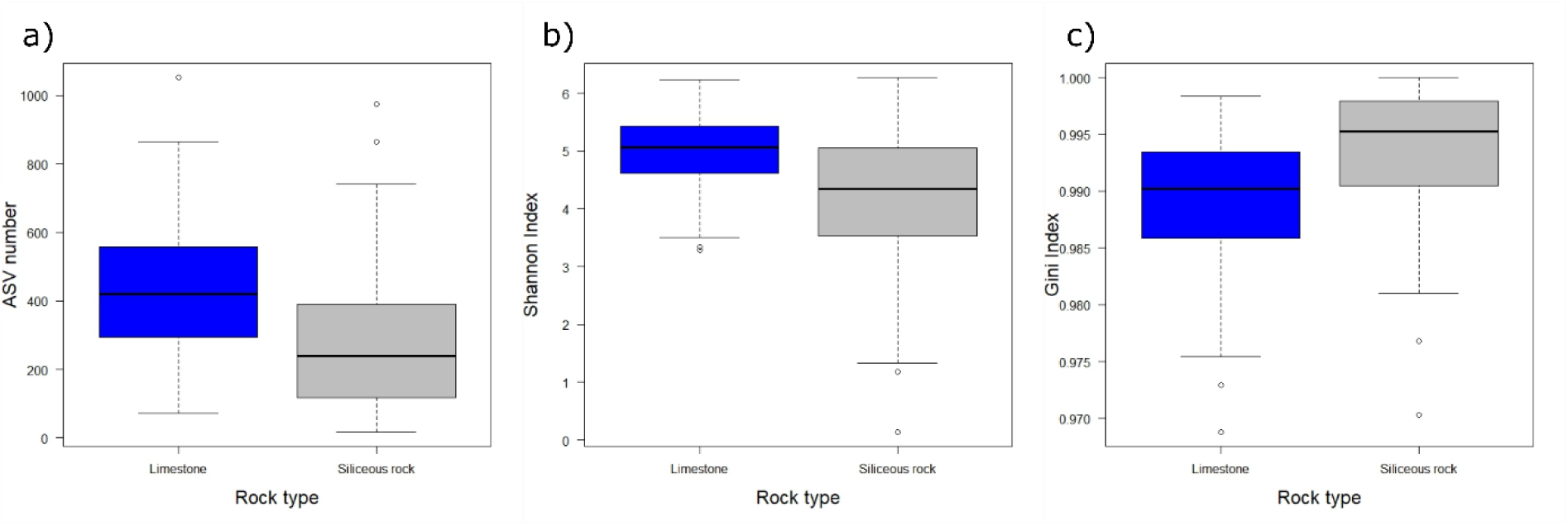
Boxplots of the Amplicon Sequences Variants number (a) Shannon index (b) and Gini index (c) in relation to the rock substrate (blue = limestone, grey = siliceous rock). The thick lines represent the median, boxes upper and lower limits the 25^th^ and the 75^th^ percentiles respectively, whiskers the data points beyond the 5^th^ percentile (lower whisker) and the 95^th^ percentile (upper whisker), open circles represent the outliers.

The GLM results of the most abundant phyla showed that only Actinobacteriota changed with the interaction between fragmentation and rock type (F_2,200_ = 7.214, P_FDR_ = 0.0108) (Fig. S3e). No phylum changed according to fragmentation (F_2,200_ ≥ 0.179, P_FDR_ ≥ 0.859), but Cyanobacteria, Actinobacteriota, Chloroflexi and Planctomycetota changed with the rock substrate (F_1,200_ ≥ 8.666, P_FDR_ ≤ 0.0103) (Fig.S3a-d). In particular, on siliceous rock there were more Cyanobacteria and Chloroflexi, while on limestone more Actinobacteriota and Planctomycetota were present.

All five most abundant phyla changed with elevation (F_1,200_ ≥ 5.549, P_FDR_ ≤ 0.0444) (Fig. S4), in particular Cyanobacteria, Chloroflexi and Planctomycetota decreased, while Proteobacteria and Actinobacteriota increased with higher elevation.

Proteobacteria was the only phylum to change according to eastness (F_1,200_ = 9.610, P_FDR_ = 0.0253), and Plantomycetota was the only one to change according to northness (F_1,200_ = 8.935, P_FDR_ = 0.0359) (Fig. S5). These two orders were more abundant at north and north-east expositions respectively.

Analyses performed on the most abundant orders’ relative abundances showed that Cyanobacteriales changed according to the fragmentation (F_2,202_ = 6.871, P_FDR_ = 0.019) (Fig. S6), as they were more abundant in contiguous *TC* and less abundant in the presence of predominantly fragmented rock.

Acetobacterales, Sphingomonadales, Solirubrobacterales, Rhizobiales, Isosphaerales, Thermomicrobiales, Gemmatales, Ktedonobacterales, Leptolyngbyales, Burkholderiales, Caulobacterales, Rhodobacterales and Rubrobacterales changed according to rock type (F_1,202_ ≥ 8.002, P_FDR_ ≤ 0.0231) (Fig. S7), in particular Acetobacterales, Ktedonobacterales and Leptolyngbyales were more abundant on siliceous rock, while Sphingomonadales, Solirubrobacterales, Rhizobiales, Isosphaerales, Cytophagales, Thermomicrobiales, Gemmatales, Burkholderiales, Caulobacterales, Rhodobacterales and Rubrobacterales were more abundant on limestone.

Solirubrobacterales was the only order to change according to the interaction between rock type and fragmentation (F_2,202_ = 8.481, P_FDR_ = 0.0183) (Fig. S8).

Cyanobacteriales, Solirubrobacterales, Rhizobiales, Gemmatales, Ktedonobacterales, Burkholderiales and Frankiales changed with elevation (F_2,202_ ≥ 11.482, P_FDR_ ≥ 0.00761) (Fig. S9), in particular Cyanobacteriales, Rhizobiales, Gemmatales and Ktedonobacterales decreased in abundance with elevation, while Solirubrobacterales, Burkholderiales and Frankiales increased in abundance at higher elevation.

In the end, Burkholderiales changed with eastness (F_1,202_ = 16.76, P_FDR_ = 0.00387) while Acetobacterales, Rhizobiales, Isosphaerales and Ktedonobacterales changed with northness (F_1,202_ ≥ 9.442, P_FDR_ ≤ 0.038) (Fig. S10).

Results about indicator species revealed that on limestone no indicator genera could be identified where both contiguous *TC* and fragmented rock surfaces were present and also not when combining data from contiguous *TC* only and those fragmented rocks. On the other hand, on siliceous rock, none of the indicator genera were detected for fragmented rock only (Tab. 2). All the seven ASVs classified as the genus *YB-42* were subsequently blasted on ncbi and they all had the best match with the strain *Chroakolemma pellucida 719*.

**Table 2.** Results of analysis for indicator genera for different levels of fragmentation and/or different rock substrates and their ecologically relevant combinations for TC. N.d. means that indicator genera were not detected.

| Fragmentation | Rock |  |
| --- | --- | --- |
|  | Limestone | Siliceous Rock |
| Contiguous TC | *Paracoccus*, *Lawsonella*, *Dolosigranulum* | *Streptococcus*, *Tepidimonas*, *Cohnella* |
| Both contiguous TC and fragmented rock surface | n.d. | JG30a-KF-32 |
| Contiguous TC and both contiguous TC and fragmented rock surface | n.d. | YB-42 |
| Fragmented rock surface | n.d. | n.d. |
| Fragmented rock surface and both contiguous TC and fragmented rock surface | n.d. | *Bradyrhizobium*, *Bryocella*, *Acidisoma*, *Acidicaldus* |
| All levels | *Rubellimicrobium*, *PMMR1*, *Oceanicella*, *MIZ36*, *Lysobacter*, *OM27 clade* | *Rubritepida* |
| All rock samples contiguous TC | *Acinetobacter*, *Sporosarcina*, *Haemophilus* |  |
| All rock samples contiguous TC and both contiguous TC with fragmented rock surface | *Herpetosiphon*, *Stenotrophomonas* |  |
| All samples | *Acidiphilium*, *Sphingomonas*, *Spirosoma*, *Aliterella*, *Fimbriiglobus*, *Solirubrobacter*, *Rubrobacter*, *Blastocatella*, *Pseudonocardia*, *Synechococcus* PCC-7502, *Aurantisolimonas*, *Rhizobacter*, *Kribbella*, *Chlorogloea* SAG-10-99, *Allorhizobium-Neorhizobium-Pararhizobium-Rhizobium* |  |

Finally, we evaluated the shared bacterial ASVs in contiguous *TC* samples dividing these samples in two big groups, limestone and siliceous rock. Results showed that 39 ASVs were shared between contiguous limestone TC samples. At genus level, 20 of these ASVs were not classified and the others resulted to be: *Acidiphilium, Psychroglaciecola, PMMR1*, *Candidatus Udaeobacter, Fimbriiglobus, Hymenobacter, Pseudonocardia, Rubrivirga, Solirubrobacter, Sphingomonas and Truepera* (Tab. S2). In siliceous rock, no shared ASVs were detected among contiguous *TC*.

## DISCUSSION

This paper describes the lithic bacterial communities of alpine areas of Switzerland with a novel focus on *Tintenstrich* communities (*TC*), using a metabarcoding approach. Overall, our results show that bacterial communities vary according to rock substrate, elevation and exposition, and also that *TC* differ depending on the rock substrates.

### *TC* host unique communities compared to other rock morphologies

The results of this study show that cyanobacteria and proteobacteria are the predominant phyla of rock bacterial communities. This result is consistent with the current limited knowledge of bacterial lithic communities and *TC* ^[4,5,23]^. Different results were reported for sandstone in Antarctica, where Proteobacteria and Actinobacteria were the most predominant phyla, while Cyanobacteria were under-represented. A second study reported communities with only one cyanobacterial phylotype^[6,35]^. In our sampling design, we excluded sandstone a priori, and our data are therefore complementary but not directly comparable. In travertine samples of the Arctic, Cyanobacteria were also under-represented^[7]^, and the authors also suggest that this may be the effect of the chemical composition of the rock. Our data can neither confirm nor reject these results because no travertine is present in our sampling region in Switzerland^[36]^.

Exposition, elevation, fragmentation and rock substrate all contributed to the variation of the bacterial communities presented herein, and *TC* host different communities than other rock morphologies. All these variables are intrinsically related to water availability on the rock, which is known to affect the microbial community. Interestingly, replicates within Weisshorn, an area which includes both limestone and siliceous rock, clustered separately along the y-axis of the CCA suggesting that the rock substrate can be a stronger variable than geographic location influencing the bacterial community. The variance explained by the CCA axes is low but significant, and may be explained by the high within-sample variability. Indeed, it has already been reported that microenvironmental characteristics drive the community composition (e.g., rock microarchitecture)^[11]^.

### *TC* on limestone show a higher diversity

In terms of alpha diversity, results showed that both richness and evenness are higher on limestone than on siliceous rock. This result is consistent with the fact that the nutrients on limestone are more limited to calcium carbonate^[37]^, therefore there is subsequently a lower competition as the result of a more oligotrophic environment. Furthermore, limestone is more subjected to bioerosion and rock weathering, and therefore it is a more ephemeral substrate where this dynamicity may obstruct the instauration of specific bacterial populations that need more time to develop^[4,38]^. As a matter of fact, the respiration of endolithic lichens and of all the other microorganisms inhabiting the rock substrate increases CO_2_ concentrations forming H_2_CO_3_ that subsequently decreases pH and promotes ions leaching^[39]^. Limestone is a particularly dense substrate, therefore bio-weathering is also a survival strategy adopted by the communities inhabiting this substrate to colonize the inner layers and avoid the epilithic environment where different stressors are stronger (e.g., UV radiation)^[4,40]^. On the other hand, siliceous rock is more resistant to rock weathering. Different material, of both mineral and biological origin, can consequently accumulate on siliceous rock surfaces forming micro-layers that can act as traps where nutrients can accumulate. Indeed, these micro-layers often present EPS that have adhesive properties, they keep the microorganisms together, promote their inclusion in the substrates and also promote retention of water and nutrients^[39]^. No differences of alpha diversity indexes were detected according to exposition or elevation.

### Unstable (limestone) and stable (silicious) rock environments host different taxa

GLMs performed at phylum level revealed that Actinobacteriota were more abundant on limestone than on siliceous rock and that they changed according to the interaction between fragmentation and rock substrate. Their relative abundance was constant on limestone, while there was a decrease related to the decrease of fragmentation on siliceous rocks. Actinobacteriota are an ubiquitous phylum that can colonize extreme environments and have already been reported in the lithic substrate^[35,41]^. Their decrease in relative abundance on siliceous rock, and especially on fragmented rock surfaces, can be the effect of the selective pressure due to the more stable environment. Not only limestone is a less stable substrate than siliceous rock, but also the presence of fragmented surface on siliceous rock implies a more stable siliceous substrate than contiguous *TC* only.

The other most abundant phyla changed with the rock substrate as well. Cyanobacteria and Chloroflexi were more abundant on siliceous rock, while Planctomycetota were more abundant on limestone. Similarly to Actinobacteriota, Planctomycetota are ubiquitous phyla inhabiting also extreme environments, and this can explain their similar trends. On the other hand, the higher relative abundances of Cyanobacteria and Chloroflexi on siliceous rock have different possible causes. First, Cyanobacteria are not only present as free-living bacteria, but also as cyano-lichen, and our molecular approach cannot distinguish between these two groups. When scratching the rock samples we collected, it was not possible to differentiate the biomass obtained from the rock between lichen and non-lichen material, also because it is often impossible to separate lichen from the rock since they can also grow as endoliths^[42]^. Therefore, DNA was extracted from the whole rock surface. A similar result was observed by Purahong et al.^[43]^, who showed that cyanobacteria diversity was negatively affected in highly disturbed areas. Chloroflexi, instead, are known to be involved in the formation of matrixes that allow the aggregation of external material and it is more likely that they form on a more stable substrate^[44]^. Abundance of Cyanobacteria, Chloroflexi and Planctomycetota also decreased with elevation, while abundance of proteobacteria and Actinobacteriota showed the opposite trend. An analogous explanation can be provided for this observation, as at higher elevation the environment is less stable in Switzerland because of more durable snow-packs and extreme conditions (e.g., higher UV radiation, lower temperatures and more freeze-thaw cycles). Considering this, only Planctomycetota shows a different trend from the one expected, but their trend has also a different pattern compared to the other phyla, which shows a high peak in abundance at 2000 m a.s.l. (Fig. 8e). Proteobacteria and Planctomycetota showed also a significant trend with eastness and northness respectively, and they both slightly decrease with higher solar expositions.

Results at order level also showed differences among rock substrates with elevation and with exposition. Burkholderiales (64.5 % of the sequences were unclassified at genus level) and Sphingomonadales (54.6 % of the sequences were classified as *Sphingomonas*) are not discussed because they represent heterogeneous taxonomic groups where it is not possible to identify specific metabolisms or ecological trends^[45,46]^.

On contiguous *TC,* the order Cyanobacteriales had a higher relative abundance, confirming that Cyanobacteria are enriched on *TC*. This order provides information about both filamentous and non-filamentous cyanobacteria (e.g *Nostoc*, *Gloeocapsa*). When looking at the Cyanobacteriales sequences in contiguous *TC* only, the majority of these sequences (60.74 %) were classified at family level as Chroococcidiopsaceae and the second most represented family is the one of Nostocaceae (11.71 % of the sequences), and these two families were the two most abundant cyanobacterial ones in these types of samples. These observations are consistent with the fact that both filamentous and non-filamentous Cyanobacteria are involved in *TC* formation, with a predominance of non-filamentous ones. Furthermore, there is no match with the results of the analysis about indicator genera, where no genera belonging to this order are reported for all contiguous TC samples. The lack of an indicator genera may suggest that there is more differentiation according to the rock type and to other environmental variables at the lowest taxonomic levels.

The order Acetobacterales also changed with rock type and exposition. Most of the sequences of this order were classified as *Acidiphilium* at genus level (48.8 %) and it was the far more abundant genus within this order (43.5 % of the sequences were unclassified). The frequent classification of *Acidiphilium* is consistent with its higher relative abundance on siliceous rock which is a more acidic substrate and, therefore, a more suitable environment for this genus^[47]^. *Acidiphilium* also decreased with solar exposition, possibly related to the lower persistence of water on sun exposed surfaces, and therefore, less bioleaching and acid rock drainage^[4]^.

Solirubrobacterales were more abundant on limestone than on siliceous rock and they also changed with the interaction between rock type and fragmentation. This order is known to be present in soil, it positively correlates with the presence of nutrients and can decrease with soil erosion^[48]^. Therefore, our results were rather unexpected, since limestone is more oligotrophic and more subjected to erosion and weathering phenomena^[4,38]^.

While most of the sequences of the order Rhizobiales were unclassified (58 %), their higher relative abundance on siliceous rock is consistent with the fact that it is a more stable environment where roots system can develop, which are known to represent a good substrate for most of the genera belonging to this order^[49]^. The tight relation with root systems may also explain the decrease of Rhizobiales with elevation.

The order Gemmatales relative abundance was higher on limestone. This order is chemoheterothrophic with an optimum pH of growth typically being neutral-acidic^[50]^. Therefore, these limestone-inhabiting Gemmatales genera may be more neutrophilic to better colonize this rock type than siliceous rock.

Ktedonobacterales were higher on siliceous rock and abundance decreased with elevation and exposition, presenting 66.17 % of their sequences unclassified at order level, while the most abundant genus was *1959-1* (32.4 %). This predominance on siliceous rock is consistent with previous discovery in a quartzite cave, where Ktedonobacterales was dominant in the first stage of orthoquartzite rock alteration^[51]^.

The order Leptolyngbyales resulted also more abundant on siliceous rock, and this result is consistent with the fact that the SILVA genus *YB-42* of this order has been recognized as an indicator cyanobacterium for *TC* on siliceous rock from our analysis. Previous results, on the other hand, show that the order Leptolyngbyales is also present in sandstone and dolomite^[8,12,20,22]^ and is already known to be composed by rock biofilm-forming bacteria^[4]^. Therefore, this order may be overall compatible with the rock environment, and possible substrate-specificity may occur at more specific taxonomic levels.

The order Caulobacterales was more abundant on limestone and most of its sequences were classified at genus level as *PMMR1* (77.6 %). While not much information is available about this order, the presence on limestone is consistent with the oligotrophic nature of caulobacterales^[52]^.

Rhodobacterales were more abundant on limestone and 82 % of the sequences were classified at genus level as *Rubellimicrobium*. This genus is present in both air and soil and is known for its metal tolerance^[53,54]^ but no additional information is available in the current literature to explain the trend in this study.

Finally, Rubrobacterales were also more present in limestone samples and all sequences were classified as *Rubrobacter.* This genus has an ability to tolerate radiation and desiccation^[55]^, but no further data are available to justify this trend.

### Indicator genera characterize different levels of fragmentation especially on limestone

Results about indicator genera revealed further differences between limestone and siliceous rock. On limestone, contiguous *TC* and all limestone samples taken together were the only two groups having indicator genera. On the contrary, on siliceous rock we had indicator genera for all the levels except for fragmented rock surface. These results may be interpreted as a higher beta diversity on siliceous rock. Interestingly, the indicator genus of *TC* on siliceous rock (both contiguous and with fragmented rock surface) resulted to be *Chroakolemma pellucida 719,* a filamentous cyanobacterium first isolated in arid soils biocrust^[56]^. Data about siliceous rocks from the Pamir mountains revealed the presence of the family Leptolyngbyaceae (the one of *Chroakolemma pellucida 719)* in three samples out of eight, underlying that this bacterium cannot be found in all siliceous rock samples^[11]^.

All contiguous *TC* of limestone samples shared 20 ASVs, while no shared ASVs were detected on contiguous *TC* of siliceous rocks. This observation supports the hypothesis that limestone has a lower beta diversity than siliceous rocks. Furthermore, limestone nutrients are limited to calcium carbonate, and therefore it is plausible that the total richness of the community is similar between sample sites since only genera with similar key functions and high adaptation can colonize such nutrient-limited environment. Furthermore, the CCA shows that all limestone samples cluster in a more restricted area (Fig. 4), and two fragmentation levels resulted not different in limestone only. Siliceous rock, on the other hand, has a more spread distribution of the samples on the CCA biplot (Fig. 4) and the community change at all fragmentation levels. This spread in the samples is also consistent with the fact that siliceous rock can contain more different minerals, allowing higher variability of the community. More detailed information of the substrate composition may allow to obtain more defined clusters.

It is important to provide more data concerning these communities with a special focus on cyanobacteria. Indeed, more than 2425 secondary metabolites specifically produced by cyanobacteria have been described so far, also in lichen symbiosis^[16,57,58]^. Some of these metabolites are recognized as cyanotoxins in the water quality guidelines of the World Health Organization^[59]^ while many other metabolites demonstrate inhibitory effects on metabolite enzymes and other bioactivities^[60]^. The semi-aquatic nature of *TC* and their connection with waters may facilitate the release of both bacteria and their metabolites in water catchments, rivers or lakes. This aspect is extremely important, especially in a not so far future scenario where glaciers will disappear, aquifers will carry less water, and occasional surface water may become essential to human and wildlife (e.g., for mountain huts). The monitoring of water quality will consequently gain more relevance, even in small catchments. While one study that investigated soil crusts, did not detect cyanotoxin-producing genes^[61]^, their presence on the rock surface cannot be excluded in general. Unfortunately, most of the studies on secondary metabolites producing cyanobacteria have been conducted in aquatic environments and no strains have been isolated from non-aquatic or semi-aquatic environments with this focus. The prevalence of cyanobacteria in *TC*, emphasizes that the presence of cyanotoxin-encoding genes and the presence of other secondary metabolites from cyanobacteria may be promising in the Swiss Alpine region.

## CONCLUSIONS

This article provides a first molecular description of *Tintenstrich* communities. Results show that cyanobacteria play an important role in forming these structures, and that alpha diversity depends on the stability and on the nutrients present in the substrate. One cyanobacterial genus only resulted to be a good indicator of *TC* on siliceous rock, but the scarcity of knowledge of these environment does not allow to obtain a high resolution at the lowest taxonomic levels. More studies should focus on such environments, since their ecological role and their composition may result soon important for water quality assessment.

## Data availability

Sequences were submitted to NCBI (https://www.ncbi.nlm.nih.gov/sra/PRJNA1095436).

## Acknowledgements

We acknowledge financial support of the Eawag-WSL collaboration “Blue-Green Biodiversity Research Initiative” to the project “Blue-Green Cyanobacteria: Diversity, Toxins and alpine Tourism” to CS, EJ and SF. CS acknowledges the High Altitude Research Station Jungfraujoch for the support in field work (linked to the project “Ecophysiology, growth and reproduction of Umbilicaria virginis, a nival lichen”).

