## Supplementary material for "Lithic bacterial communities: ecological aspects focusing on *Tintenstrich communities*"

Francesca Pittino[1,2,3*], Sabine Fink[1], Juliana Oliveira[1,2], Elisabeth M.-L. Janssen[2], Christoph Scheidegger[1]

[1]*Biodiversity and Conservation Biology, Swiss Federal Research Institute (WSL), Birmensdorf, Switzerland.*

[2]*Department of Environmental Chemistry, Swiss Federal Institute of Aquatic Science and Technology (EAWAG), Dübendorf, Switzerland.*

[3]*Department of Earth and Environmental Sciences, University of Milano-Bicocca, Milan, Italy*

**Supplementary material**

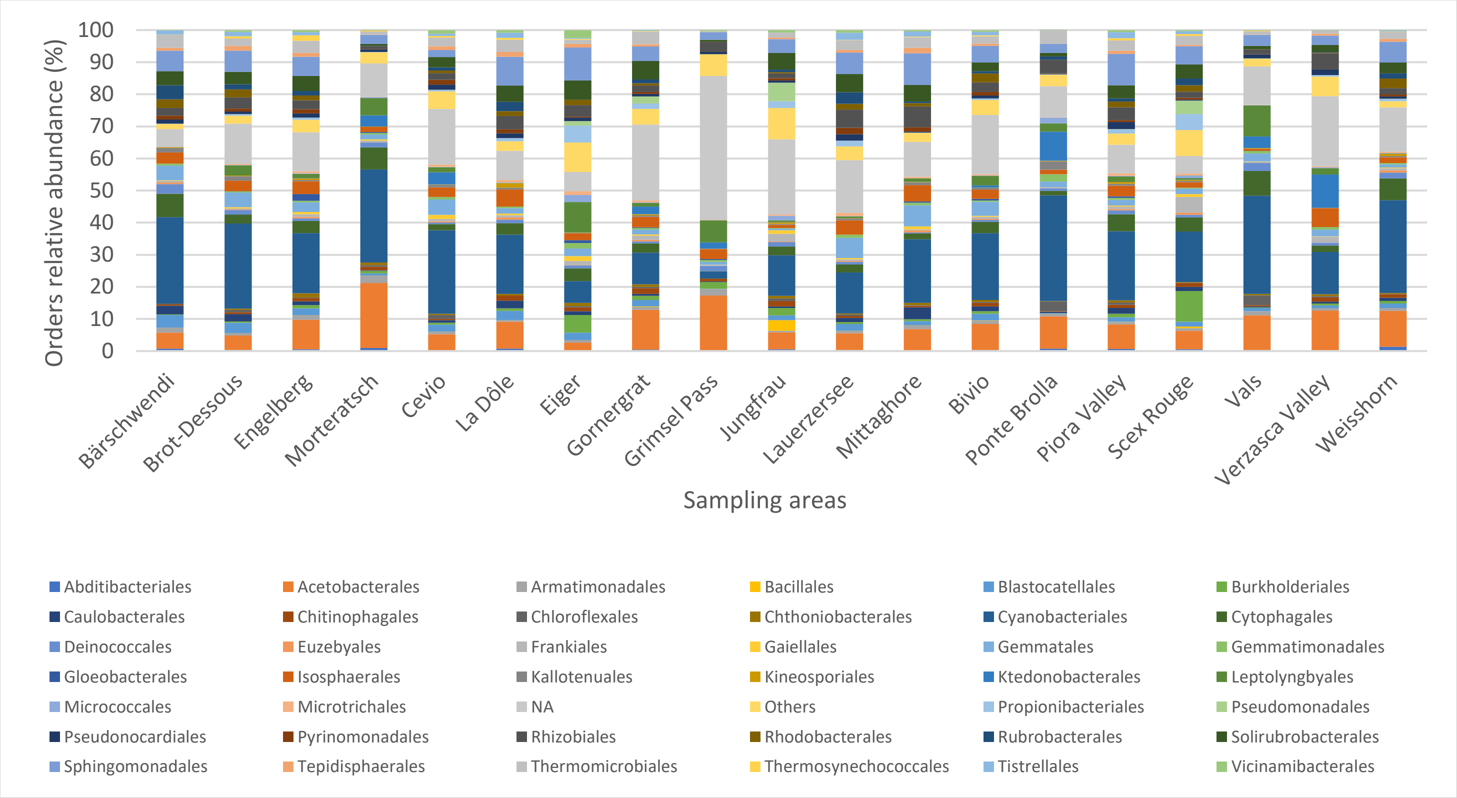


*Figure S1 Relative abundance of bacterial orders expressed as the percentage of sequences. Only the most abundant orders are shown, those which are not included between the most abundant were grouped in 'Others'.*

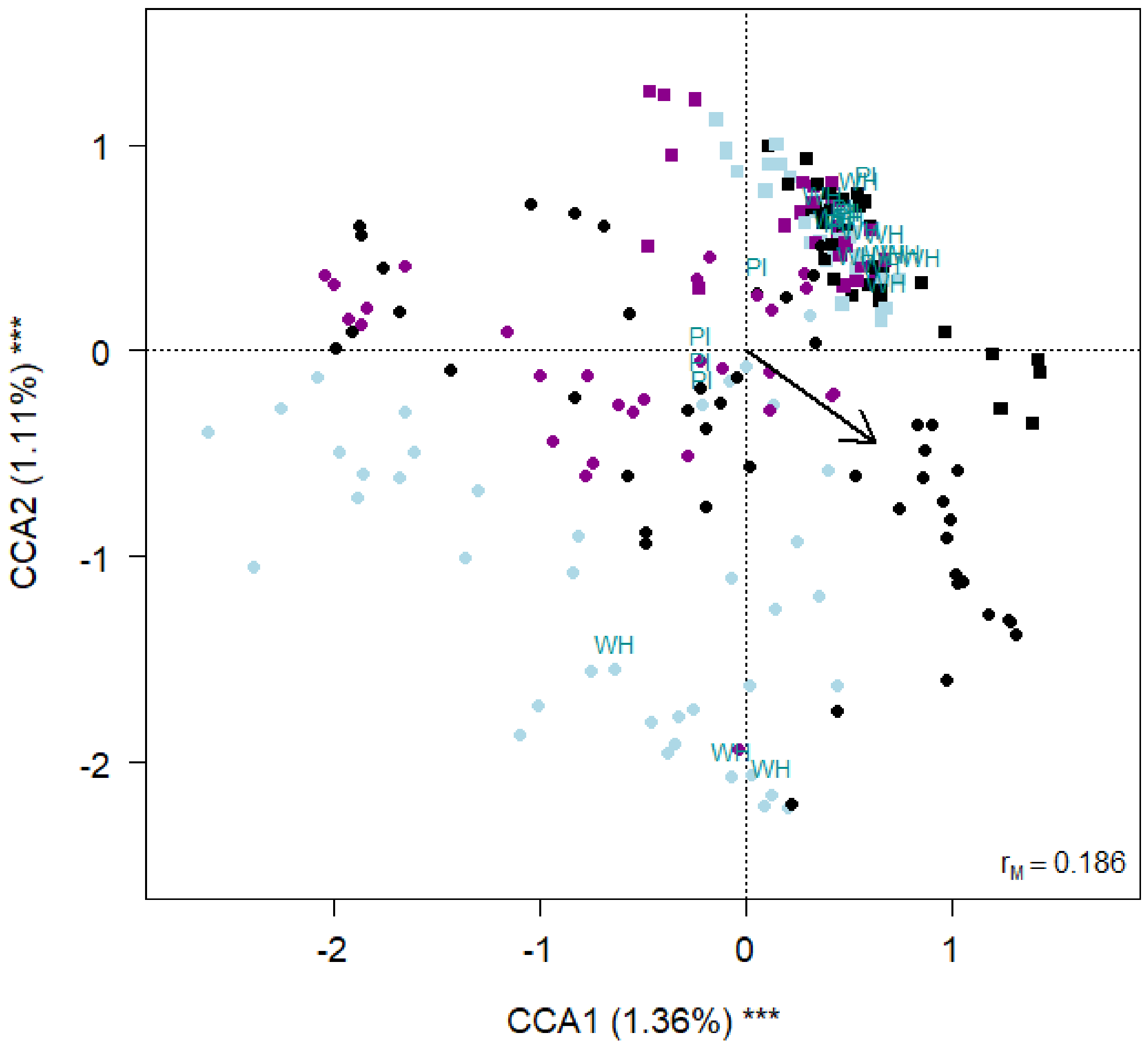


*Figure S2 Biplot from the CCA on bacterial ASV abundance on smoothness, rock type, elevation, northness and eastness. Each point represents one sample. The smoothness is indicated by different colours (black = smooth tintenstrich, purple = Tintenstrich with other material, light blue = Mixed material). The arrow indicates the elevation. Squares indicate limestone samples and circles siliceous rock samples. A few sampling areas are reported: WH = Wheisshorn, PI = Piora Valley. The percentage of variance explained by each axis and its significance (***: P< 0.001) is reported. rM is the Mantel correlation coefficient between the chi-square distance between samples and the Euclidean distance between the corresponding symbols in the graph. Values close to one indicate that the graph correctly represents the distance between samples.*

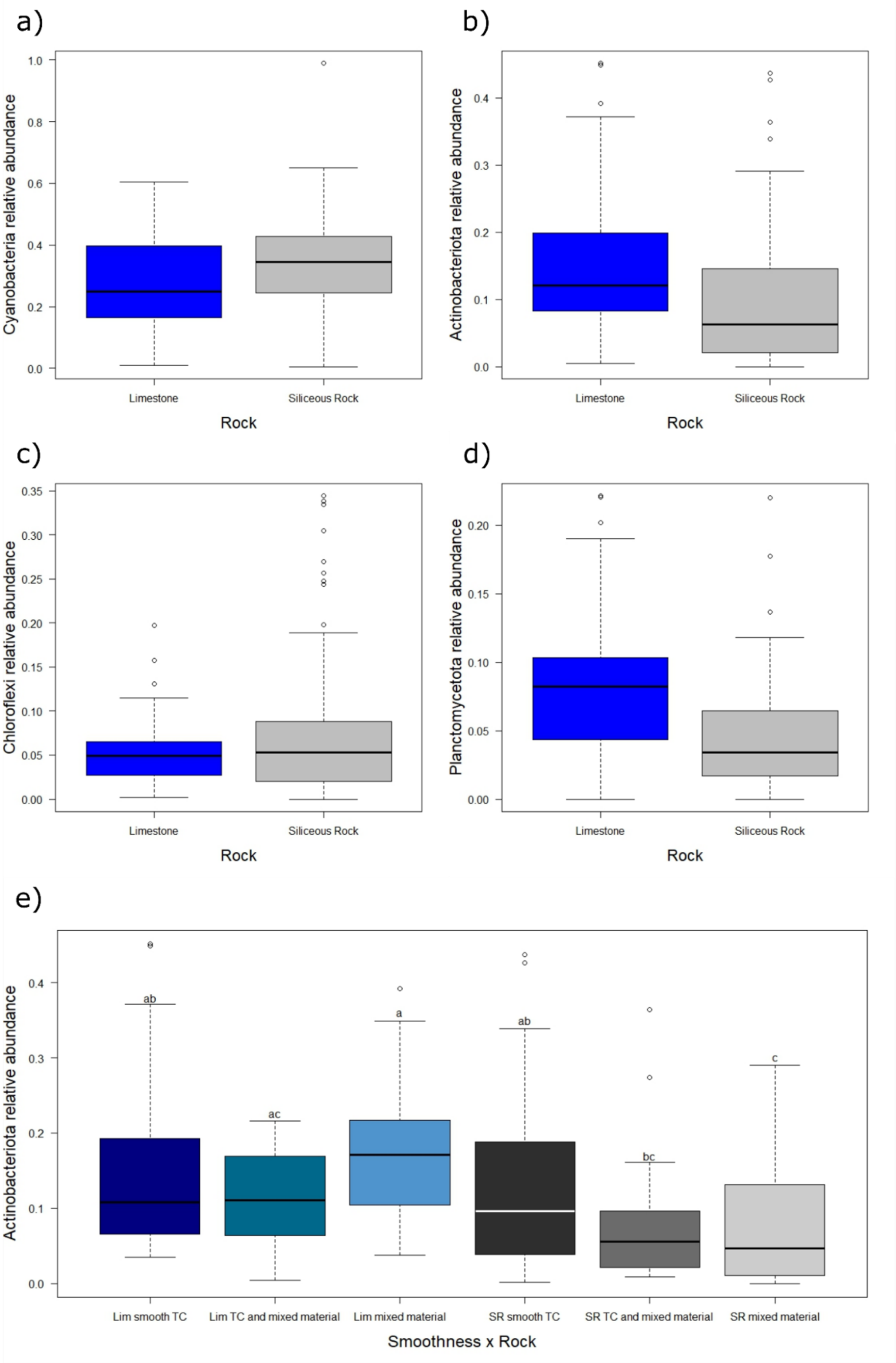


*Figure S3. Boxplots of the relative abundances of Cyanobacteria (a), Actinobacteriota (b), Chloroflexi (c) and Planctomycetota (d) according to the rock substrates: limestone (blue) and siliceous rock (grey). The thick lines represent the median, boxes upper and lower limits the 25$^{th}$ and the 75$^{th}$ percentiles respectively, whiskers the data that go beyond the 5$^{th}$ percentile (lower whisker) and the 95$^{th}$ percentile (upper whisker), open circles represent the outliers and different letters indicate differences between the mean values of different groups.*

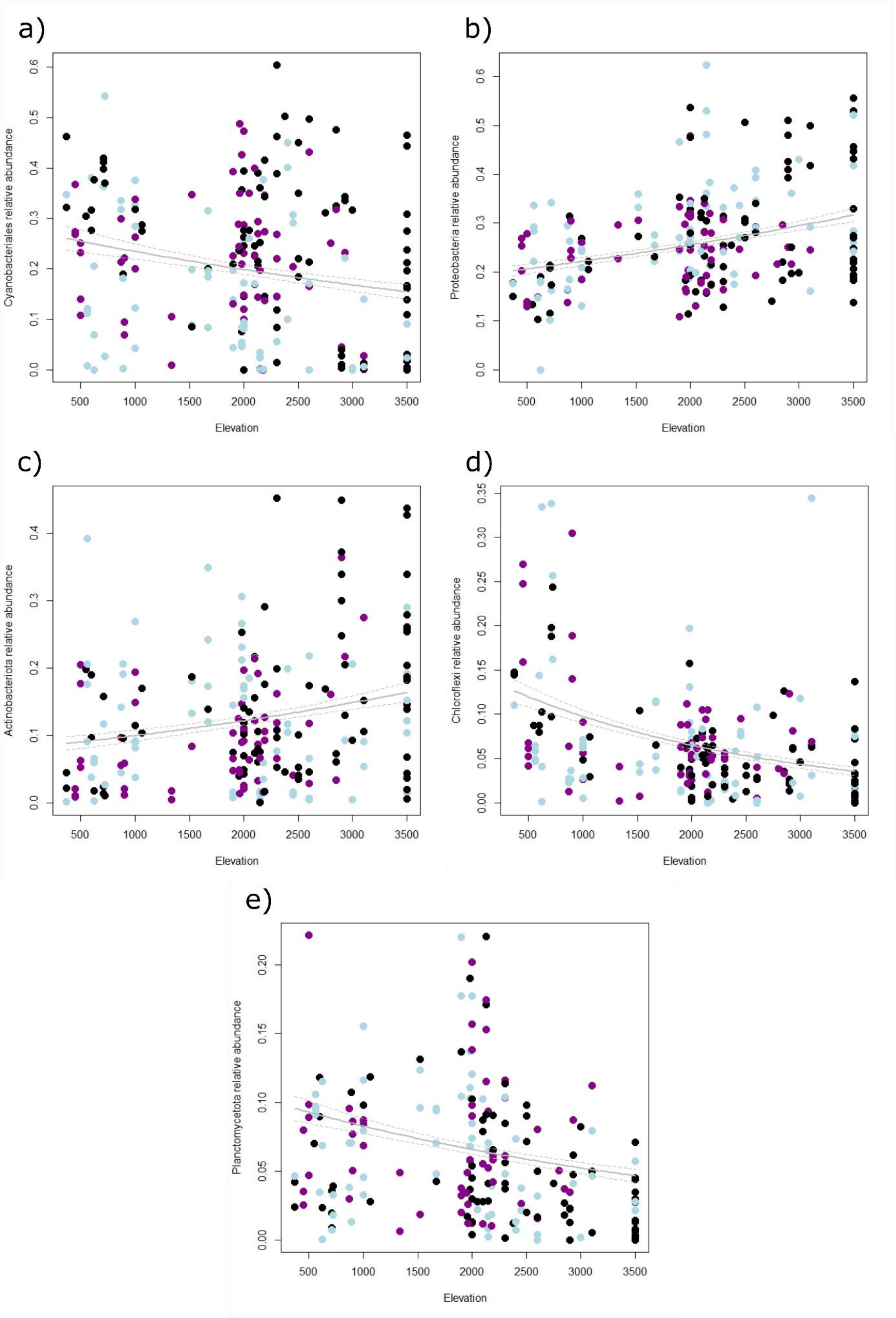
a)
Cyanobacteriales relative abundance
0.0
0.1
0.2
0.3
0.4
0.5
0.6
500
1000
1500
2000
2500
3000
3500
Elevation
b)
Proteobacteria relative abundance
0.0
0.1
0.2
0.3
0.4
0.5
0.6
500
1000
1500
2000
2500
3000
3500
Elevation
c)
Actinobacteriota relative abundance
0.0
0.1
0.2
0.3
0.4
500
1000
1500
2000
2500
3000
3500
Elevation
d)
Chloroflexi relative abundance
0.00
0.05
0.10
0.15
0.20
0.25
0.30
0.35
500
1000
1500
2000
2500
3000
3500
Elevation
e)
Planctomycetota relative abundance
0.00
0.05
0.10
0.15
0.20
500
1000
1500
2000
2500
3000
3500
Elevation

*Figure S4 GLM plots showing the variation of cyanobacteria (a), proteobacteria (b), actinobacteriota (c), chloroflexi (d) and planctomycetota (e) with elevation. The smoothness is indicated by different colours (black = smooth tintenstrich, purple = Tintenstrich with other material, light blue = Mixed material).*

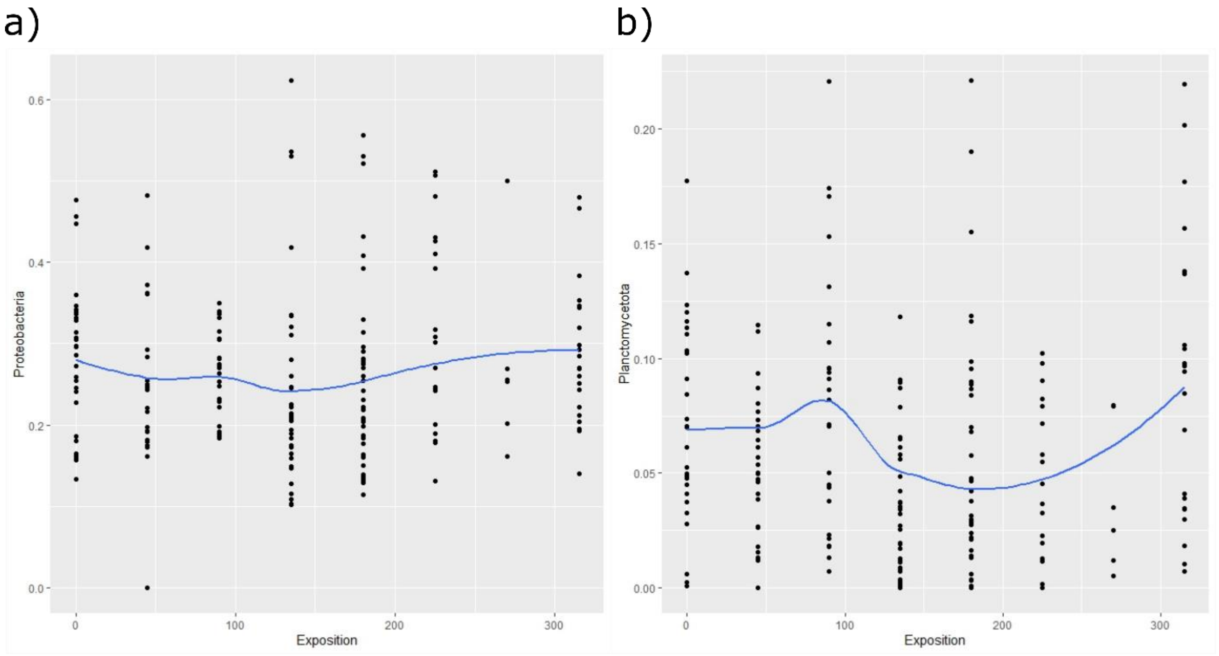


*Figure S5 GLM biplot showing the variation of proteobacteria (a) and planctomycetota (b) according to exposition.*

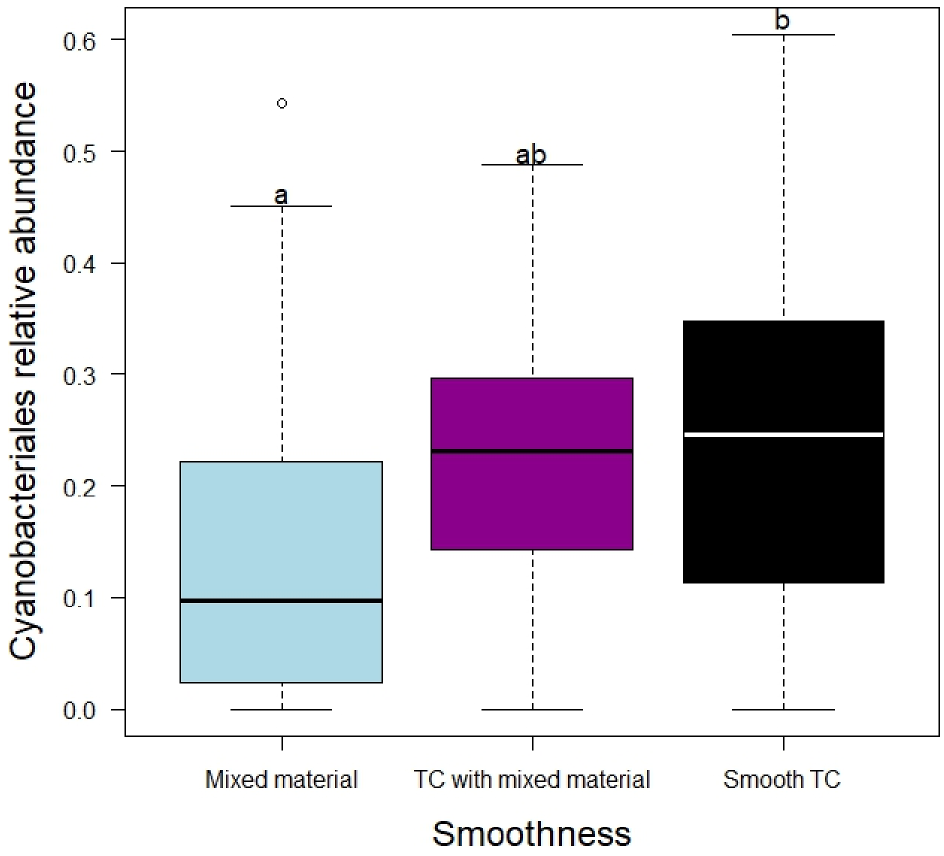


*Figure S6 Boxplots of the relative abundances of cyanobacteriales in different levels of rock smoothness (mixed The thick lines represent the median, boxes upper and lower limits the 25th and the 75th percentiles respectively, whiskers the data that go beyond the 5th percentile (lower whisker) and the 95th percentile (upper whisker), dots represent the outliers and different letters indicate differences between the mean values of different groups.*

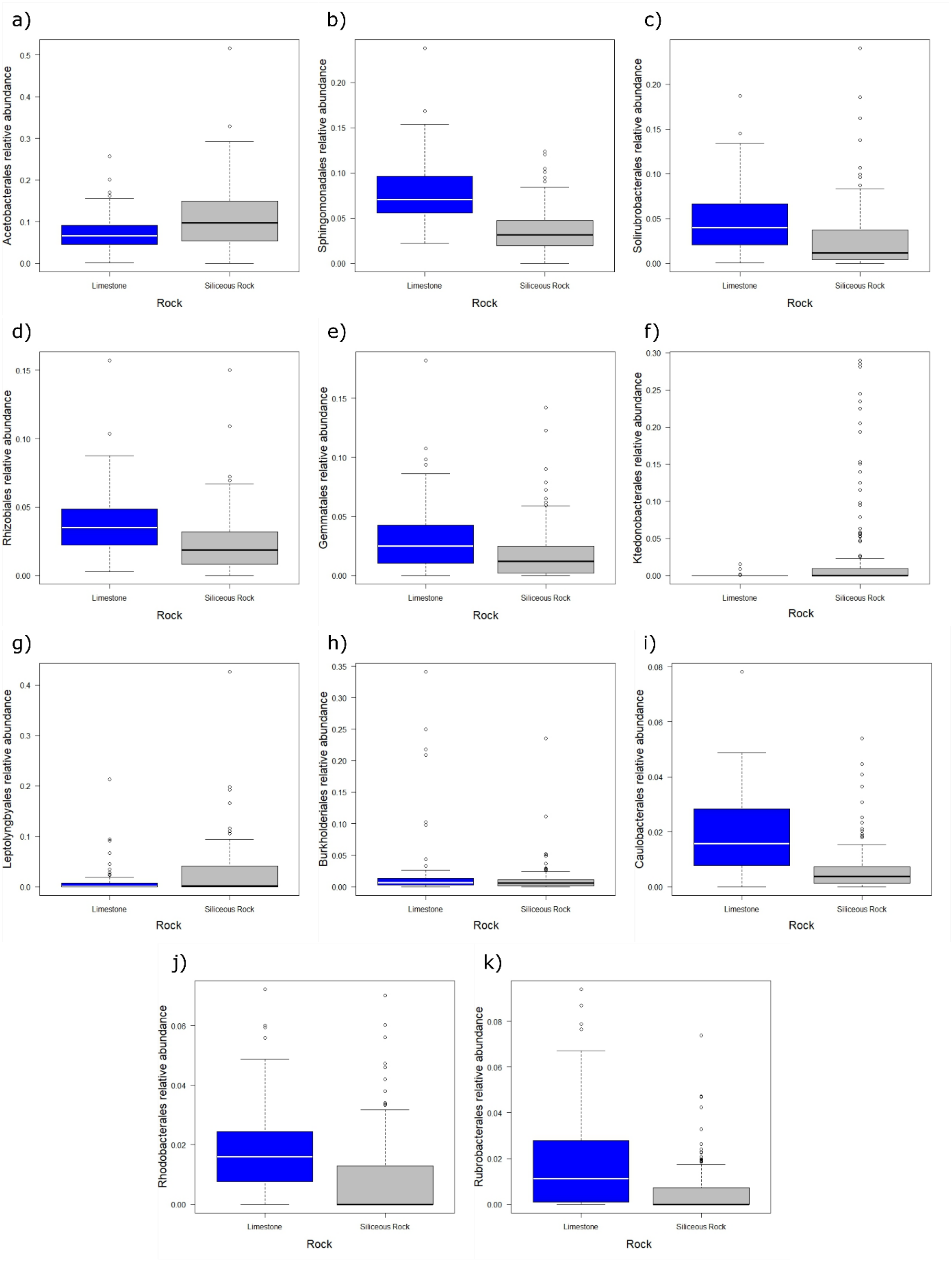


*Figure S7 Boxplots of the relative abundances of acetobacterales (a), sphingomonadales (b), solirubrobacterales (c), rhizobiales (d), gemmatales (e), ktedonobacterales (f), leptolyngbyales (g), burkholderiales (h), caulobacterales (i), rhodobacterales (j) and rubrobacterales (k) showing differences according to the rock substrate (blue = limestone, grey = siliceous rock). The thick lines represent the median, boxes upper and lower limits the 25$^{th}$ and the 75$^{th}$ percentiles respectively, whiskers the data that go beyond the 5$^{th}$ percentile (lower whisker) and the 95$^{th}$ percentile (upper whisker), dots represent the outliers and different letters indicate differences between the mean values of different groups.*

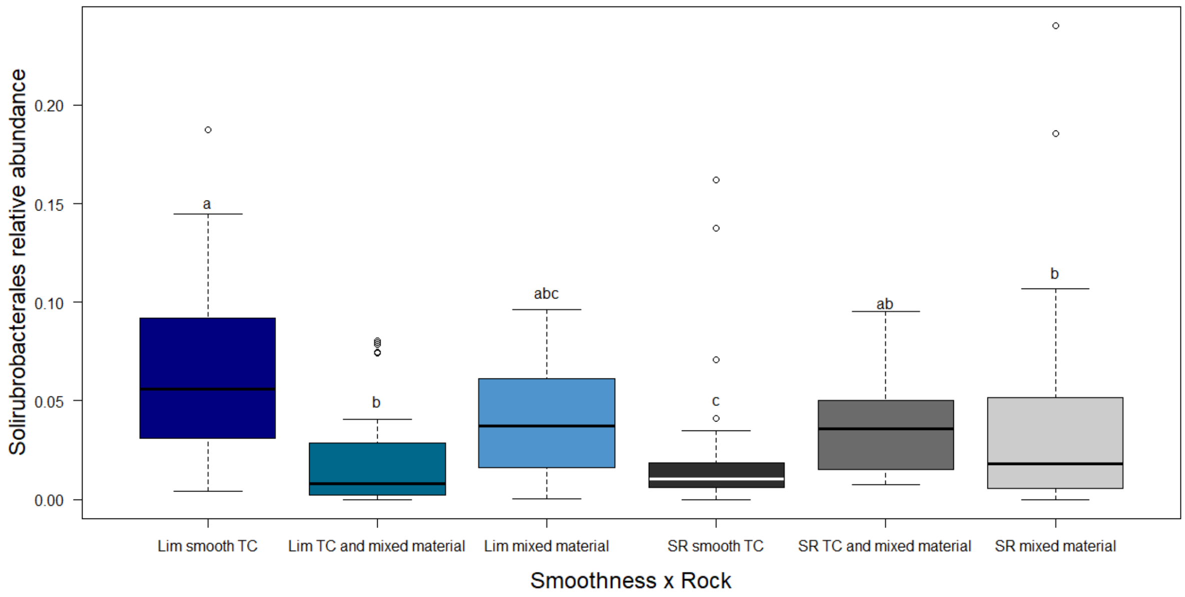


*Figure S8 Boxplots of the relative abundances of solirubrobacterales on the interaction between smoothness and rock substrate (Lim = limestone in blue shades, SR = Siliceous Rock in grey shades). The thick lines represent the median, boxes upper and lower limits the 25th and the 75th percentiles respectively, whiskers the data that go beyond the 5th percentile (lower whisker) and the 95th percentile (upper whisker), dots represent the outliers and different letters indicate differences between the mean values of different groups*

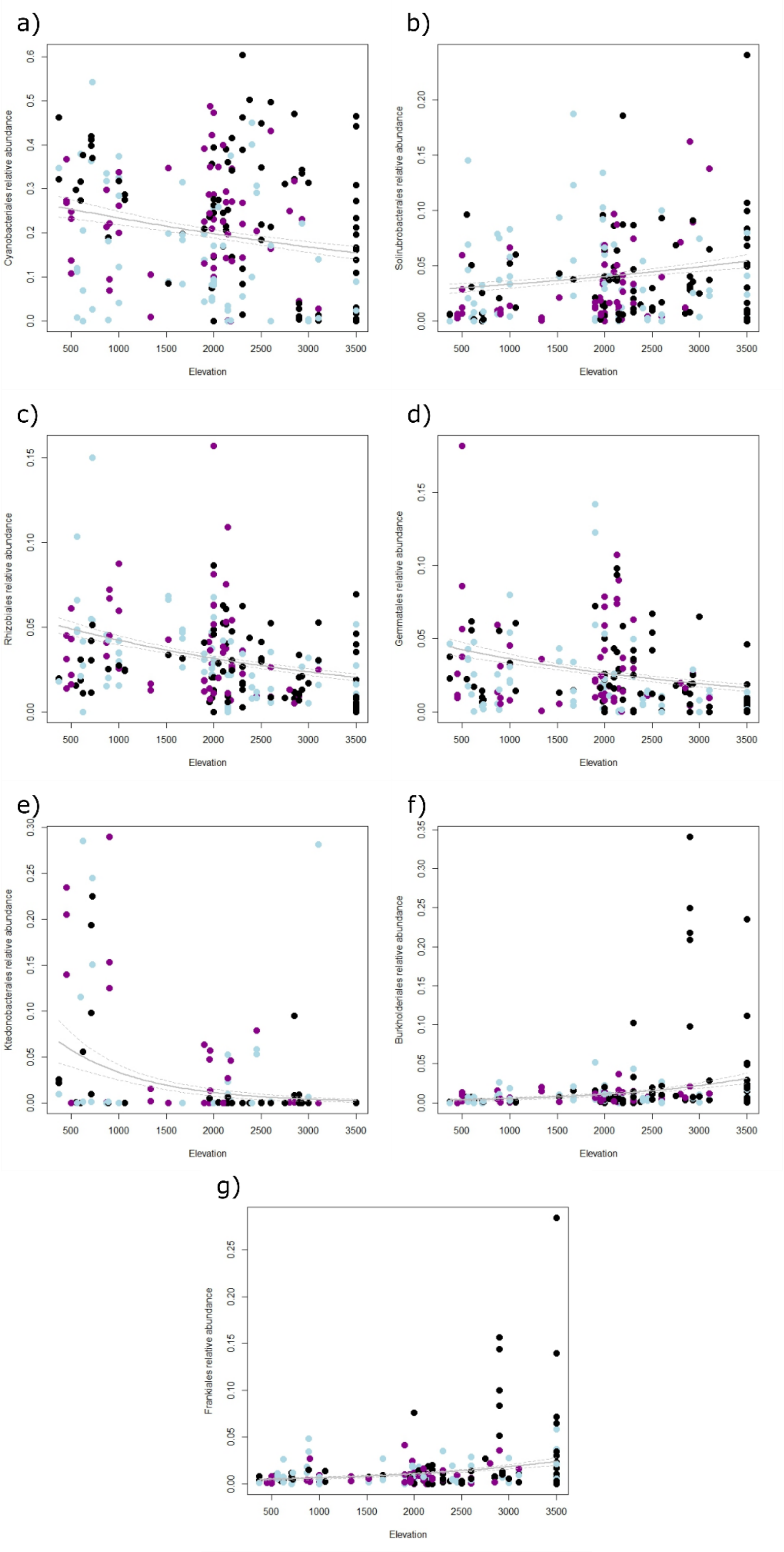


*Figure S9 GLM biplots showing the trend of Cyanobacteriales (a), Solirubrobacterales (b), Rhizobiales (c), Gemmatales (d), Ktedonobacterales (e), Burkholderiales (f) and Frankiales (g) according to elevation.*

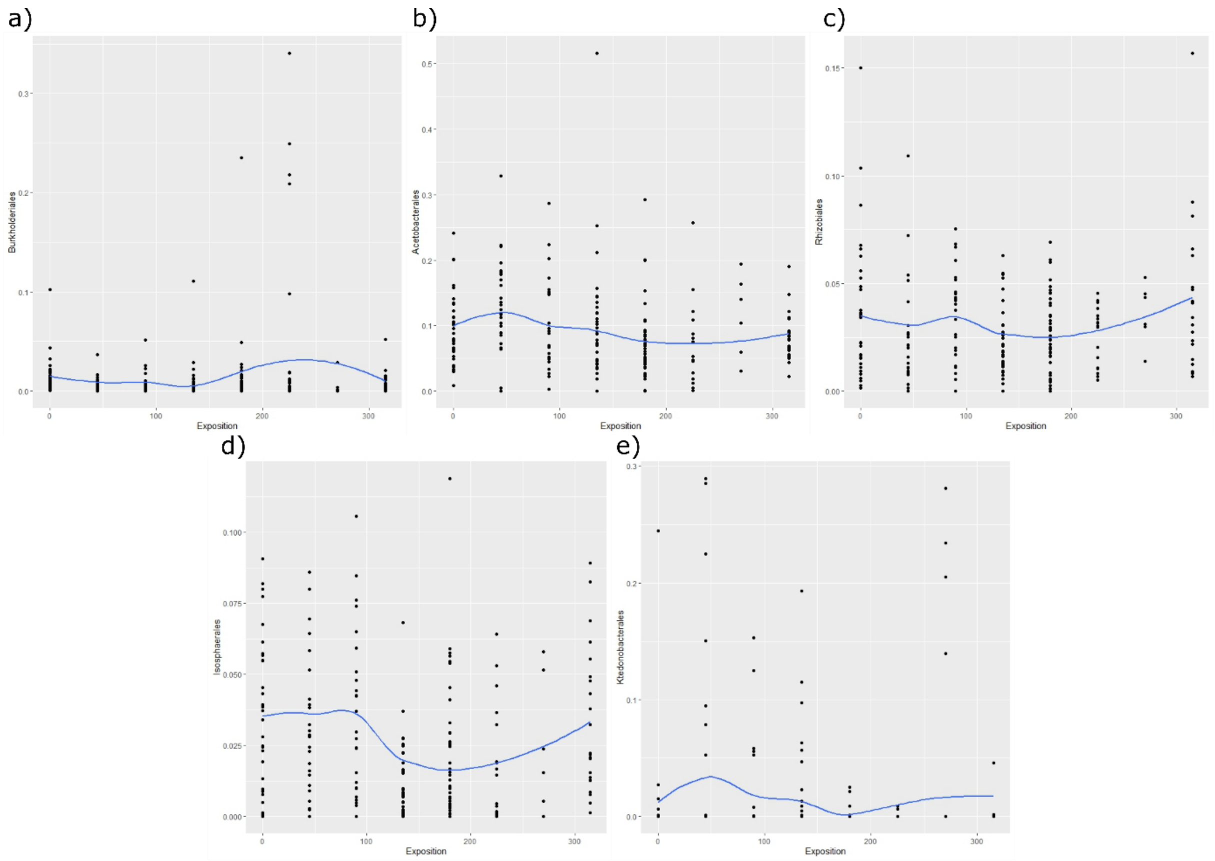


*Figure S10 GLM biplots showing the trend of burkholderiales (a), acetobacterales(b), Rhizobiales (c) and Isosphaerales (d) and Ktedonobacterales (e) according to exposition. Exposition is expressed as Azimuth degrees.*

*Table S1 List of samples and their respective coordinates.*

| Sample.Name | Lon | Lat |
|---|---|---|
| B1 | 8,14 | 46,7635212 |
| B2 | 8,142718341 | 46,7635212 |
| S2 | 8,4178825 | 46,7892273 |
| T1 | 8,48626 | 46,79226 |
| T2 | 8,4897 | 46,79289 |
| J_1_1_21 | 7,98554 | 46,54751 |

| | | |
|---|---|---|
| J_1_2_21 | 7,98554 | 46,54751 |
| J_1_3_21 | 7,98554 | 46,54751 |
| J_1_4_21 | 7,98554 | 46,54751 |
| J_2_1_21 | 7,98632 | 46,54776 |
| J_2_2_21 | 7,98636 | 46,54739 |
| J_2_3_21 | 7,98636 | 46,54739 |
| J_3_1_21 | 7,98636 | 46,55264 |
| J_3_2_21 | 7,98636 | 46,55264 |
| J_4_1_21 | 7,98636 | 46,55264 |
| J_4_2_21 | 8,0026 | 46,55257 |
| J_4_3_21 | 8,0026 | 46,55257 |
| J_4_4_21 | 8,00246 | 46,55272 |
| J_4_5_21 | 8,00246 | 46,55272 |
| J_4_6_21 | 8,00246 | 46,55272 |
| J_4_7_21 | 8,00246 | 46,55272 |
| J_5_1_21 | 8,00251 | 46,55271 |
| J_5_2_21 | 8,00251 | 46,55271 |
| J_5_3_21 | 8,00251 | 46,55271 |
| J_5_4_21 | 8,00251 | 46,55271 |
| J_6_1_21 | 7,97096 | 46,54527 |

| | | |
|---|---|---|
| J_6_2_21 | 7,97096 | 46,54527 |
| J_6_3_21 | 7,97096 | 46,54527 |
| J_6_4_21 | 7,97096 | 46,54527 |
| J_6_5_21 | 7,97096 | 46,54527 |
| EIG_1_1_21 | 7,97609 | 46,57592 |
| EIG_1_2_21 | 7,97609 | 46,57592 |
| EIG_1_3_21 | 7,97609 | 46,57592 |
| LW_1_1_21 | 8,61666 | 47,03758 |
| LW_1_2_21 | 8,616666667 | 47,0375 |
| LW_1_3_21 | 8,616388889 | 47,0375 |
| LW_1_4_21 | 8,616388889 | 47,0375 |
| LW_2_1_21 | 8,61956 | 47,02858 |
| LW_2_2_21 | 8,61954 | 47,02863 |
| LW_2_3_21 | 8,61969 | 47,02865 |
| LW_2_4_21 | 8,61943 | 46,02864 |
| BD_1_1_21 | 6,72805 | 46,95949 |
| BD_2_1_21 | 6,72776 | 46,95932 |
| BD_2_2_21 | 6,72776 | 46,95932 |
| BD_2_3_21 | 6,72776 | 46,95932 |
| BD_5_1_21 | 6,72757 | 46,95934 |

| | | |
|---|---|---|
| BD_4_1_21 | 6,727524 | 46,95932 |
| BD_4_2_21 | 6,727524 | 46,95932 |
| BD_3_2_21 | 6,726374 | 46,96083 |
| BD_3_3_21 | 6,726383 | 46,96083 |
| DL_1_1_21 | 6,10319 | 46,42234 |
| DL_1_2_21 | 6,10312 | 46,42236 |
| DL_1_3_21 | 6,10312 | 46,42236 |
| DL_2_1_21 | 6,09973 | 46,42408 |
| DL_2_2_21 | 6,09973 | 46,42408 |
| DL_2_3_21 | 6,09973 | 46,42408 |
| DL_4_1_21 | 6,09788 | 46,42284 |
| DL_4_2_21 | 6,09802 | 46,42288 |
| DL_4_3_21 | 6,09789 | 46,42285 |
| GR_1_1_21 | 8,30986 | 46,58922 |
| GR_1_2_21 | 8,30986 | 46,58923 |
| GR_1_3_21 | 8,30986 | 46,58924 |
| GR_2_1_21 | 8,31006 | 46,58901 |
| GR_2_2_21 | 8,31007 | 46,58902 |
| GR_2_3_21 | 8,31008 | 46,58903 |
| SUP_1_1_21 | 9,63471 | 46,46427 |

| | | |
|---|---|---|
| SUP_1_2_21 | 9,63479 | 46,4643 |
| SUP_2_1_21 | 9,63498 | 46,4643 |
| SUP_2_2_21 | 9,63498 | 46,4643 |
| SUP_3_1_21 | 9,63603 | 46,46484 |
| SUP_3_2_21 | 9,63603 | 46,46484 |
| SUP_4_1_21 | 9,64239 | 46,46763 |
| SUP_4_2_21 | 6,72642 | 46,96088 |
| SUP_4_3_21 | 6,72642 | 46,96000 |
| NE_1_1_21 | 9,6565 | 46,479 |
| NE_1_2_21 | 9,6567 | 46,48003 |
| NE_1_3_21 | 9,65629 | 46,4804 |
| NE_2_1_21 | 9,65788 | 46,47069 |
| NE_2_2_21 | 9,65788 | 46,47069 |
| NE_2_3_21 | 9,65627 | 46,46991 |
| NE_2_4_21 | 9,65478 | 46,46929 |
| NE_2_5_21 | 9,65424 | 46,46921 |
| NE_2_6_21 | 9,65424 | 46,46921 |
| NE_2_7_21 | 9,65424 | 46,46921 |
| BV_1_1_21 | 9,91628 | 46,41798 |
| BV_1_2_21 | 9,91643 | 46,41785 |

| | | |
|---|---|---|
| BV_1_3_21 | 9,9162 | 46,41778 |
| BV_2_1_21 | 9,92647 | 46,41677 |
| BV_2_2_21 | 9,92613 | 46,41669 |
| BV_2_3_21 | 9,92675 | 46,41714 |
| SR_1_1_21 | 7,22128 | 46,31342 |
| SR_1_2_21 | 7,22128 | 46,31342 |
| SR_1_3_21 | 7,22128 | 46,31342 |
| SR_1_4_21 | 7,22128 | 46,31342 |
| SR_1_5_21 | 7,22128 | 46,31342 |
| SR_2_1_21 | 7,22451 | 46,30959 |
| SR_2_2_21 | 7,22451 | 46,30959 |
| SR_2_3_21 | 7,22451 | 46,30959 |
| SR_2_4_21 | 7,22514 | 46,30956 |
| MT_1_1_21 | 7,27459 | 46,36579 |
| MT_1_2_21 | 7,27459 | 46,36579 |
| MT_1_3_21 | 7,27392 | 46,36604 |
| MT_1_4_21 | 7,27415 | 46,36614 |
| MT_2_1_21 | 7,27374 | 46,36478 |
| MT_2_2_21 | 6,72642 | 46,96087 |
| MT_2_3_21 | 7,27137 | 46,36386 |

| | | |
|---|---|---|
| MT_2_4_21 | 7,27137 | 46,36386 |
| MT_2_5_21 | 7,27146 | 46,36385 |
| MT_2_6_21 | 7,27146 | 46,36385 |
| MT_2_7_21 | 7,27146 | 46,36385 |
| MT_2_8_21 | 6,72642 | 46,96085 |
| TT_1_1_21 | 8,41947 | 46,78387 |
| TT_1_2_21 | 8,41947 | 46,78387 |
| TT_1_3_21 | 8,41947 | 46,78387 |
| TT_1_4_21 | 8,41947 | 46,78387 |
| TT_2_1_21 | 8,41431 | 46,78379 |
| TT_2_2_21 | 8,41426 | 46,78379 |
| TT_2_3_21 | 8,41426 | 46,78379 |
| TT_2_4_21 | 8,41426 | 46,78379 |
| TT_3_1_21 | 8,40682 | 46,79024 |
| TT_3_2_21 | 8,40682 | 46,79024 |
| TT_3_3_21 | 8,40682 | 46,79024 |
| BR_1_1_21 | 8,40991 | 46,84389 |
| BR_1_2_21 | 8,40991 | 46,84389 |
| BR_1_3_21 | 8,40991 | 46,84389 |
| BR_1_4_21 | 8,40966 | 46,84389 |

| | | |
|---|---|---|
| BR_1_5_21 | 8,40966 | 46,84389 |
| BR_1_6_21 | 8,40961 | 46,84397 |
| BR_1_7_21 | 8,40961 | 46,84397 |
| BR_1_8_21 | 8,40961 | 46,84397 |
| GOR_1_1_21 | 7,78057 | 45,98266 |
| GOR_1_2_21 | 7,78057 | 45,98266 |
| GOR_1_3_21 | 7,77863 | 45,98251 |
| GOR_1_4_21 | 7,77801 | 45,98257 |
| GOR_1_5_21 | 7,77801 | 45,98257 |
| GOR_1_6_21 | 7,77447 | 45,984 |
| GOR_1_7_21 | 7,77389 | 45,98415 |
| GOR_1_8_21 | 7,76807 | 45,98366 |
| GOR_1_9_21 | 7,76693 | 45,98389 |
| GOR_1_10_21 | 7,76038 | 45,98253 |
| GOR_1_11_21 | 7,75648 | 45,98541 |
| PI_1_1_21 | 8,72076 | 46,54692 |
| PI_1_2_21 | 8,72063 | 46,54704 |
| PI_1_3_21 | 8,72049 | 46,5471 |
| PI_1_4_21 | 8,72023 | 46,54684 |
| PI_1_5_21 | 8,71907 | 46,5473 |

| | | |
|---|---|---|
| PI_1_6_21 | 8,71774 | 46,54738 |
| PI_2_1_21 | 8,71642 | 46,55481 |
| PI_2_2_21 | 8,7167 | 46,5549 |
| PI_2_3_21 | 8,7168 | 46,55489 |
| PI_2_4_21 | 8,71708 | 46,555 |
| WH_1_1_21 | 9,63781 | 46,79058 |
| WH_1_2_21 | 9,64771 | 46,7906 |
| WH_1_3_21 | 9,64779 | 46,79065 |
| WH_1_4_21 | 9,64776 | 46,79075 |
| WH_2_1_21 | 9,64667 | 46,79091 |
| WH_2_2_21 | 9,64667 | 46,79091 |
| WH_2_3_21 | 9,6465 | 46,79269 |
| WH_3_1_21 | 9,6391900 | 46,79091 |
| WH_3_2_21 | 9,6391900 | 46,79092 |
| WH_3_3_21 | 9,6392247 | 46,7909049 |
| WH_3_4_21 | 9,63918 | 46,79093 |
| WH_3_5_21 | 9,63847 | 46,79066 |
| WH_4_1_21 | 9,63759 | 46,78912 |
| WH_4_2_21 | 9,63763 | 46,78909 |
| WH_4_3_21 | 9,63921 | 46,79091 |

| | | |
|---|---|---|
| WH_4_4_21 | 9,63781 | 46,78909 |
| PE_1_1_21 | 9,94178 | 46,44186 |
| PE_1_2_21 | 9,94178 | 46,44186 |
| PE_1_3_21 | 9,94178 | 46,44186 |
| PE_1_4_21 | 9,94178 | 46,44186 |
| PE_2_1_21 | 9,943 | 46,43561 |
| PE_2_2_21 | 9,94285 | 46,43566 |
| PE_2_3_21 | 9,94273 | 46,43551 |
| PB_1_1_21 | 8,74207 | 46,19276 |
| PB_1_2_21 | 8,742089 | 46,19276 |
| PB_1_3_21 | 8,742057 | 46,19276 |
| PB_2_1_21 | 8,74166 | 46,19367 |
| PB_2_2_21 | 8,74154 | 46,19363 |
| PB_2_3_21 | 8,74154 | 46,19361 |
| PB_3_1_21 | 8,75089 | 46,18858 |
| PB_3_2_21 | 8,750903 | 46,18858 |
| PB_3_3_21 | 8,750916 | 46,18858 |
| PB_4_1_21 | 8,76016 | 46,18869 |
| PB_4_2_21 | 8,76009 | 46,18842 |
| PB_4_3_21 | 8,76011 | 46,18844 |

| | | |
|---|---|---|
| VZ_1_1_21 | 8,78492 | 46,29759 |
| VZ_1_2_21 | 8,7851 | 46,29755 |
| VZ_1_3_21 | 8,78488 | 46,29762 |
| VZ_2_1_21 | 8,79059 | 46,2999 |
| VZ_2_2_21 | 8,79054 | 46,30004 |
| VZ_2_3_21 | 8,79054 | 46,30004 |
| VZ_2_4_21 | 8,79054 | 46,30004 |
| VZ_3_1_21 | 8,79239 | 46,2833 |
| VZ_3_2_21 | 8,79198 | 46,28362 |
| VZ_3_3_21 | 8,79205 | 46,28366 |
| CV_1_1_21 | 8,59607 | 46,31686 |
| CV_1_2_21 | 8,59607 | 46,31686 |
| CV_1_3_21 | 8,59629 | 46,31687 |
| CV_2_1_21 | 8,5963 | 46,31736 |
| CV_3_1_21 | 8,60091 | 46,32122 |
| CV_4_1_21 | 8,59669 | 46,3104 |
| CV_4_2_21 | 8,59669 | 46,3104 |
| CV_4_3_21 | 8,59669 | 46,3104 |
| VL_1_1_22 | 9,11462 | 46,58091 |
| VL_1_2_22 | 9,11473 | 46,58087 |

| | | |
|---|---|---|
| VL_1_3_22 | 9,11465 | 46,58098 |
| VL_1_4_22 | 9,11475 | 46,58094 |
